# A scalable, low-cost phenotyping strategy to assess tuber size, shape, and the colorimetric features of tuber skin and flesh in potato breeding populations

**DOI:** 10.1101/2023.08.14.553050

**Authors:** Max J. Feldman, Jaebum Park, Nathan Miller, Collins Wakholi, Katelyn Greene, Arash Abbasi, Devin A. Rippner, Duroy Navarre, Cari Schmitz Carley, Laura M. Shannon, Rich Novy

**Affiliations:** Temperate Tree Fruit and Vegetable Research Unit, USDA – Agricultural Research Service, Prosser, WA; Small Grains and Potato Germplasm Research Unit, USDA – Agricultural Research Service, Aberdeen, ID; Department of Botany, University of Wisconsin-Madison, Madison, WI; Horticultural Crops Production and Genetic Improvement Research Unit, USDA – Agricultural Research Service, Prosser, WA; The Beacom College of Computer and Cyber Sciences, Dakota State University, Madison, SD; Aardevo North America, LLC, Boise, ID; Department of Horticultural Sciences, University of Minnesota, St. Paul, MN

**Keywords:** Potato, *Solanum tuberosum*, phenotype, machine vision, heritability, machine learning, hollow heart

## Abstract

Tuber size, shape, colorimetric characteristics, and defect susceptibility are all factors that influence the acceptance of new potato cultivars. Despite the importance of these characteristics, our understanding of their inheritance is substantially limited by our inability to precisely measure these features quantitatively on the scale needed to evaluate breeding populations. To alleviate this bottleneck, we developed a low-cost, semi-automated workflow to capture data and measure each of these characteristics using machine vision. This workflow was applied to assess the phenotypic variation present within 189 F1 progeny of the A08241 breeding population. Our results provide an example of quantitative measurements acquired using machine vision methods that are reliable, heritable, and can be used to understand and select upon multiple traits simultaneously in structured potato breeding populations.

## Introduction

The merits of the potato (*Solanum tuberosum*), particularly its yield potential and nutritional benefits relative to other crops, have made it an essential global staple food along with rice and wheat (Hirsch et al., 2013). The U.S. is a major producer of potatoes (∼ 21 million tonnes in 2019), a majority of which are used to manufacture processed foods (67%), particularly French fries and potato chips (∼78%) (USDA National Agricultural Statistics Service, 2020). Consumer preference for these processed food products have necessitated increased breeder focus on tuber sugar and acrylamide levels, texture, specific gravity, and tuber shape to consistently maintain the quality requirements of their finished products (Fajardo et al., 2013). Tuber size and shape are especially important characteristics because they determine compatibility for use in processing machines including peelers, cutters, and fryers, etc. as well as the portion of a usable raw product after cutting and trimming (Bradeen & Kole, 2016; Pavek & Knowles, 2015; Van Eck et al., 1994; Tao et al., 1995). Unlike grain crops, genetically identical potato clones can exhibit substantial variation in tuber size and shape across field locations and growth years, making it difficult to standardize the size and shape of any single potato clone (Lindqvist-Kreuze et al., 2015; Van Eck et al., 1994). As such measurement of tuber size, shape, and the variance of these characteristics is an essential component of both the breeding and clone evaluation process.

Tuber size is traditionally quantified by gravimetric, mechanical, or optical sorting methods and used to partition large tuber lots into categorical subsets of similar size. Tuber size determination within large, bulk harvested tuber lots from late-stage trials (tens of clones, > 15 hill plot) frequently relies upon the acquisition and maintenance of dedicated sorting equipment. Estimating tuber size in studies performed on early generation breeding material (hundreds of clones, < 10 hill) is often performed manually using calipers (Lindqvist-Kreuze et al., 2015; Prashar & Jones, 2014; Van Eck et al., 1994) or by separating individual tubers into categorical descriptors by hand (Alsahlany et al., 2021; Bradshaw et al., 2008; Buhrig et al., 2015; Domański, 2001; Hara-Skrzypiec et al., 2018; Li et al., 2005; Lindqvist-Kreuze et al., 2015; Śliwka et al., 2008).

Description of tuber shape can be quantified either through ordinal numbering evaluation (Alsahlany et al., 2021; Bradshaw et al., 2008; Domański, 2001; Hara-Skrzypiec et al., 2018;. Li et al., 2005; Lindqvist-Kreuze et al., 2015; Śliwka et al., 2008; USDA Plant Variety Protection Office, 2015) or as the relationship between individual measurements (Lindqvist-Kreuze et al., 2015; Prashar & Jones, 2014; Tabatabaeefar, 2002; Van Eck et al., 1994). Visual scoring of tuber shape into ordinal categories is a rapid, quantitative way to assess shape. Categorical descriptors of shape range in resolution from 2 shapes (round or long) all the way up to 6 shape categories (long, long oval, oval, round-oval, round, compressed). These types of values are relatively easy for humans to interpret but are limited in their descriptive capacity due to poor measurement resolution (ordinal categories do not reflect the continuous variation in tuber shape), differences in the magnitude of shape variance between tuber populations (all russet type tuber lot vs. primitive introgression population), and variation introduced due to human bias (discordance between human potato graders) (Poland & Nelson, 2011).

Length – width ratio (L/W ratio), also known as aspect ratio, is a commonly used, quantitative tuber shape descriptor. Aspect ratio greater than 1.75 is an important tuber characteristic that can determine the suitability of a clone or tuber lot for French fry production. This descriptor readily discriminates between chipping and russet clones. These high-resolution measurements can be acquired inexpensively using calipers or a ruler and be easily interpreted between experiments performed on different tuber population types. Tabulation of tuber length, width, and their relationship provide greater depth and content than ordinal shape values. Unfortunately they require greater effort to compile in terms of labor and recording on a per tuber basis, limiting application within both basic research and mass production systems of the food industry. Additional shape characteristics, such as tapered ends, or tuber bend is not typically quantified in breeding populations.

Colorimetric characteristics of tuber skin and flesh are also key features that determine the suitability of potato cultivars for different fresh market and processing market classes. These features are important descriptors that may be used to predict pathogen infection or the presence of tuber blemishes. Much of our knowledge pertaining to the genetic basis of potato skin and flesh color has been ascertained from manual qualitative scoring of individual clones for their color characteristics (Salaman, 1910; Zhang et al., 2009), whereas visual inspection of skin quality and internal flesh color is an important component of tuber grading protocols. To date, most research studies have relied upon categorical scoring of skin and flesh color performed by human subjects (Buhrig et al., 2015; De Jong, 1987; Jung et al., 2009) or based upon subsampling within a single tuber using a digital colorimeter (Buhrig et al., 2015; Qin et al., 2020; Roe et al., 2014; Thornton et al., 2013; Waterer, 2010). These types of measurement strategies are subject to many of the challenges associated with manual measurements of tuber shape; particularly the tradeoffs observed between accuracy, precision, resolution, and throughput (Krupek et al., 2021; Miller et al., 2023; Roe et al., 2014).

The propensity of a clone to develop tuber defects is another important characteristic that must be considered during the potato cultivar development process. Certain defects including susceptibility to hollow heart, spraing, tuber greening, in-field sprouting, growth cracks and knobby tuber outgrowth (secondary growth) are unacceptable and clones expressing these defects must be identified and removed from breeding programs; whereas other defects including tuber bruising, skinning, and scab or nematode susceptibility are important but can sometimes be addressed through agronomic management strategies. The presence of hollow heart is somewhat common and relatively easy to identify, making it an excellent test case to evaluate the feasibility of machine learning based classification. This physiological, internal tuber defect that can occur when growth in the perimedullary region of the tuber outpaces growth of the pith causing the development of irregularly shaped cavities within the tuber pith tissue (Rex & Mazza, 1989). Frequently, expression of this defect is a consequence of water or nutrient fluctuation during the growth season. Genetics has been demonstrated to contribute to hollow heart susceptibility and in some cases susceptibility is likely independent of final tuber size (Jansky & Thompson, 1990). The invasive and low-throughput methods needed to evaluate internal defects prevent rigorous evaluation during early generation selection process and limit our ability to understand the genetic basis of these characteristics. Fortunately, work towards the detection and quantification of tuber defects is becoming more prevalent in the literature (Marino et al., 2019; Miller et al., 2023; Moallem et al., 2013; Wang & Xiao, 2021).

Application of machine vision to quantify tuber size, shape, and color has been under development by numerous groups since the late 1980s (J. Marchant et al., 1988; J. A. Marchant et al., 1990; McClure & Morrow, 1987; Miller et al., 2023; Neilson et al., 2021). Data acquisition using digital imaging and machine vision is an inexpensive, high-throughput technique that can capture accurate measurements of multiple continuous traits simultaneously (Li et al., 2014). Additionally, image data is long lived and can be revisited later for further analysis. Multiple research groups report high correlation between manual and machine vision-based measurement of tuber size (Hasankhani & Navid, 2012; Neilson et al., 2021; Noordam et al., 2000; Si et al., 2017; Zhou et al., 1998), tuber shape (Hasankhani & Navid, 2012; Liu et al., 2021; Miller et al., 2023; Neilson et al., 2021; Noordam et al., 2000; Si et al., 2017; Su et al., 2017; Tao et al., 1995; Zhou et al., 1998), or categorical descriptors of tuber color (Barnes et al., 2009; Caraza-Harter & Endelman, 2020; Miller et al., 2023). Generally, this process involves assigning pixels within a digital image that encompass the potatoes using either manual thresholding (Miller et al., 2023; Neilson et al., 2021; Si et al., 2017; Zhou et al., 1998) or machine learning techniques (Noordam et al., 2000; Si et al., 2017; Tao et al., 1995) and calculating measurements from of the shape contour of each tuber (Miller et al., 2023; Neilson et al., 2021; Noordam et al., 2000; Si et al., 2017; Tao et al., 1995; Zhou et al., 1998). In some cases, these measurement strategies have been applied to collect and analyze data at the throughput needed in mechanical tuber grading platforms (Hasankhani & Navid, 2012; Noordam et al., 2000; Rady & Guyer, 2015; Si et al., 2017; Zhou et al., 1998).

Surprisingly, outside of breeding value estimation for parental clones (Neilson et al., 2021), this technology has not been widely applied in the field of quantitative genetics in potato. The routine linkage and association mapping techniques have been deployed in many other cropping systems require quantitative measurement of plant phenotypes for each clone within breeding populations that consist of hundreds-to-thousands of individual clones (Bernardo, 2019). We speculate this is largely because without mechanization, data collection is laborious and the mechanical data collection platforms deployed require expensive, dedicated use equipment that is not well suited for the evaluation of early generation breeding trials (hundreds of clones, < 10 hills/plot) at small-to-medium size research stations. Knowledge derived from such studies can be applied to better understand the underlying genetic architecture of these important traits and be applied in marker assisted/genomic selection breeding schema.

In this study we utilize an image-based potato tuber measurement system, based on consumer grade sensors and open-source data analysis workflows constructed with beginner friendly, high-level programming languages (Python, R). Using student labor, we applied this workflow to measure tuber size, tuber shape, colorimetric characteristics of tuber skin and flesh, and hollow heart susceptibility on 189 F1 progeny within the A08241 breeding population, replicated twice, using several different techniques. Our results indicate that quantitative measurements acquired using this scalable methodology are reliable and can be used to measure, understand, and select upon multiple traits simultaneously in potato breeding populations.

## Materials and Methods

### Plant material

The autotetraploid mapping population (A08241), comprised of 189 F1 progeny, was obtained from a cross between a cultivar ‘Palisade Russet’ (female parent) and a breeding clone ‘ND028673B-2Russ’ (male parent). This mapping population was initially created for late blight resistance study and has been grown and maintained at USDA-ARS Small Grains and Potato Germplasm Research Unit in Aberdeen, ID. For this study, the two parents and their progeny were planted in a randomized complete block design with two replications of eight-hill plots in 2019. Both Palisade Russet (Novy et al., 2012) and ND028673_2Russ (Dr. Susie Thompson, North Dakota State University, personal communication) show oblong or long tuber shape. For all the experiments in this study we measured ten tubers from each clone (five tubers from replication 1 and another five tubers from replication 2).

### Digital caliper measurement

After harvesting, five tubers per clone replicate (ten total tubers per clone) were randomly selected and then their length and width were measured by a digital caliper. The length was defined as the distance between stem-end and bud-end of the tuber. The definition of width was the longest distance perpendicular to the length. The ratio of tuber length divided by tuber width is reported as aspect ratio [aka length / width ratio (L/W ratio)]. The units for both distance measurements are expressed in centimeters.

### SolCAP visual assessment for tuber shape and weight measurement

Tuber shape of the whole population was visually scored from “1” (Compressed) to “5” (Long) (Park et al., 2021). This measurement is called “SolCAP Visual Assessment (SVA)” in this research project. For SVA, five tubers from each replication of each clone were horizontally aligned on a plate first. After observing the five tubers, a researcher assigned a single score representative of the entire sample, which reflects the average tuber shape of the five tubers. The weight [unit: ounces (*oz*)] of each individual was also measured together. After completing data collection, the SVA and weight data were compared with the digital caliper measurement. A comparison between replication 1 and 2 data was also conducted.

### Image acquisition

Digital images of up to 5 whole tubers per replicate were acquired of the same five tubers/replicate that were measured previously with the digital caliper. A Nikon D7100 DSLR camera with a Nikon AF-S DX NIKKOR 18-140mm f/3.5-5.6G ED VR Lens (focal length 18 mm) was suspended 32” above the floor of the imaging cabinet using a K&F Concept TM2534T DSLR Camera Tripod (66”) and counterbalanced with a 1-gallon water container was used for digital imaging. Tubers were labeled with a permanent marker and arrayed in numerical order under full power LED illumination (color temperature 5500K) within a HAVOX HPB-80D photo studio on both a non-reflective black background and a Laboratory Supplies CO., Inc. Cat. G129 Model A Illuminator equipped with two Philips 30-Watt Linear T12 Fluorescent Tube Light Bulbs (color temperature 6500K). Two faces of each tuber were imaged on both the black background and the illuminator light box by acquiring an image of each tuber side (take image, then rotate tuber and take another image) on the black background (with the illuminator turned off) and repeating this procedure again on top of the illuminator with the bulbs powered on. The image acquisition process and camera settings were configured using a custom Python script run on a Raspberry Pi 3 computer that minimized the potential for labeling mix ups and standardized the camera settings across all images. Optimal camera settings were determined by autofocus detection algorithms on the camera, recorded, and applied automatically to each image capture using the Python *gphoto2* library. When imaging on the black background images were captured with white balance set to 5560K, ISO equal to 100, a F-Stop value of 7.1, and a shutter speed of 1/200 seconds. On the illuminator box, white balance and ISO were kept at 5560K and 100 respectively, whereas F-Stop was increased to 8 and shutter speed was reduced to 1/250 seconds. The resulting JPEG images were written to the Raspberry Pi 3 computer and also saved on the camera’s SD card.

Digital scans of tuber flesh were acquired using a Hewlett Packard HP ColorJet 6200C flatbed digital scanner controlled by the *sane* library in a custom Python program run on a Raspberry Pi 3 computer. Before scanning each tuber was cut in half lengthwise and an image of the labeled side of each tuber was captured by arraying the tubers in numerical order on the scanning surface and covering the scanner with an opaque Rubbermaid storage bin to exclude ambient light.

Both size and radiometric calibration standards were included in each image. A blue plastic poker chip (37 mm diameter) was used as a size standard in all images and scans. An X-Rite ColorChecker Classic was used as a radiometric standard for the top-down imaging on the black background whereas a X-Rite ColorChecker Mini was used as color standards for the top-down imaging on the illuminator and the scanner. Python scripts used to control the imaging hardware can be found at: https://github.com/maxjfeldman/Feldman_hardware_control

### Image processing

Input files were JPEG images (4000 × 6000 pixels from the top-down camera, and 2550 × 3507 pixels from the flatbed scanner) of the potato tubers that had been arrayed in numerical order and arranged so that no two tubers were touching. Processing was achieved using custom Python scripts that rely heavily on the functionality provided by the OpenCV library. In some cases functions from the PlantCV library were also used (Fahlgren et al., 2015; Gehan et al., 2017). Briefly, images were cropped to remove unwanted background features and gaussian blur was used to obscure salt-and-pepper noise within the image. Potato tubers were isolated from background by applying threshold selection on the saturation channel within the HSV color space and b* channel in the CIE LAB color space. The binary image result after applying a logical AND set operation on both thresholded binary images provided repeatable isolation of the tuber from the image background. Contours of each individual tuber and size marker were then used to extract shape and colorimetric measurements that were subsequently written to a .csv file.

Measurement of potato tuber eccentricity was performed using the *contourMoments* function in the OpenCV library on the derived tuber contour. Eccentricity is a measurement of the of the shortest length of paths from a given vertex to reach any other vertex of the connected graph. It can be considered an measure of roundness, where 0 is a perfect circle and an ellipse exhibits a value greater than zero but less than one.

Tuber biomass profiles were calculated as reported by (Turner et al., 2018). Biomass profiles are an example of latent traits that capture structural information about an object that require biological interpretation through dimensional reduction usually using Principal component analysis (PCA), autoencoders, persistent homology barcodes, or elliptical Fourier descriptors. Each tuber contour was rotated to align the longest axis of a bounding ellipse to the vertical axis of the image, then the image was cropped to its minimal bounding rectangle and converted to a binary image. This rectangular image was then padded along the x-axis with zeros to maintain axis ratio after resizing, and the resulting square binary image was resized to a 100 x 100 pixel image. For each of the 100 rows the number of tuber pixels across the sweep was calculated and written to a .csv file. PCA was used to perform dimensional reduction before trait evaluation.

Color correction of each image was performed in MATLAB using the *colorChecker* function to identify the X-Rite ColorChecker Classic and the *measureColor* function was used to derive and apply the color correction matrix. Colorimetric features were calculated as the mean reflectance value (between 0 – 255) within each tuber contour, across each of the R, G, and B channels of the digital image. Dimensional reduction using principal component analysis was applied to these three mean color channel values (R, G, and B) using the *prcomp* function in R and the resulting data analyzed.

Python and R computer scripts used to perform this analysis can be found at: https://github.com/maxjfeldman/Park_Feldman_Tuber_Imaging_2023

### Heritability calculation

Broad-sense heritability was calculated by fitting a linear model with genotype as the only explanatory variable using the *lmer* function in the R package *lme4*. In cases where both sides of each tuber were imaged, only the data from side one (as annotated in data file) was used to fit the model. The model was fit using the following formula:

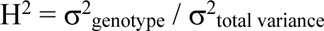

### Measurement of repeatability between tuber sides/faces

To assess measurement repeatability, values of each trait collected through machine vision techniques from top-down images of intact tubers were compared between tuber sides/faces using Pearson’s correlation coefficient. Calculations were made using the *cor* function in base R. Because only a single face of each halved tuber was collected using the flatbed scanner it was not possible to assess repeatability of tuber flesh color.

### Sample size variance calculations

To evaluate the ideal number of tuber samples needed per clone, we calculated the mean standard error of each trait using 3 to 10 tuber samples. Smaller samples were simulated from complete trait data of the 10 tubers by selecting smaller tuber samples at random. For each trait, and each clone, the standard error of each sample size was calculated using the *std_error* function from the *plotrix* package in R and then averaged across all clones. These average values of standard error were plotted to evaluate gain of precision from each additional tuber measured.

### Capture and augmentation of hollow heart dataset

Halved, individual tubers were labeled by their position in the scanner image and manually scored for the presence or absence of internal defects according to the USDA grading standards visual aid chart. This study utilized a dataset of 1500 RGB images of sliced potatoes with dimensions of 1900×1900 pixels. Out of these, only 49 images depicted potatoes with the hollow-heart defect, representing less than 4% of the total dataset (Fig. S1). The dataset is imbalanced, with a higher number of normal potato images compared to hollow-heart defect images. Imbalanced data can negatively impact the performance of a classification model, potentially leading to biased results and low accuracy. To address this, we utilized random augmentation, as described below, on the hollow-heart defect images to artificially increase the size of the dataset (Johnson & Khoshgoftaar, 2019).

Image augmentation is a technique that creates new training examples from existing ones by applying random transformations to the images. These modifications, which may include cropping, adjusting brightness and contrast, rotating, warping, and flipping, are intended to increase the size and diversity of the training dataset. Image augmentation can help to improve the performance and generalizability of machine learning models, particularly when the original dataset is small or lacks variety. In this study, a python library Albumentations (Buslaev et al., 2020) was used to perform random image augmentation on the hollow-heart potato images. The Albumentations library offers a range of transformation options, including flipping, adjusting brightness and contrast, distorting the image using a grid, rotating, scaling, and applying affine transforms. These transformations were randomly applied to each hollow-heart potato image (Fig. S2). The dataset now includes a total of 2431 images, with the number of hollow-heart defect images having been increased from 49 to 980 using augmentation.

Before image augmentation, we used a random split method to assign 80% of the images to the training set and 20% to the test set. This ensures that the model is trained on a representative sample of the data and evaluated on unseen data.

After image augmentation, the original dataset was reshaped into 3 different resolutions (256×256, 512×512, and 1024×1024) using bilinear interpolation. Deep learning models were then trained and evaluated on these reshaped datasets to determine the minimum resolution necessary for accurately classifying potatoes with hollow-heart defects from normal potatoes.

### Deep learning network architecture and network training

A convolutional neural network (CNN) was implemented for the classification of potatoes with or without the hollow-heart defect. The network (Fig. S3) contained an input layer and five blocks (B0–B5), each consisting of a 2D convolution, batch normalization, and 2D Max-pooling layer. The convolution layer utilized a 3×3 filter, zero padding to maintain the same input tensor size, and a Rectified Linear Unit (ReLU) activation function. The Max-pooling layer employed a 2×2 pool size and a stride of 2. The output of block B5 was flattened and passed through fully connected layers FC1, FC2, FC3, then the output layer (outputs 2 classes). The fully connected layers output 1024, 128, 16, and 2 values, respectively, and utilized the ReLU activation function. The CNN model was implemented using the TensorFlow deep learning framework (Abadi et al., 2016) in Python.

To train the classification CNN, the dataset that was randomly split into three parts: a training set, a validation set, and a test set. The ratio used was 6:3:1. The training set was used to optimize the model’s parameters, while the validation set was used to evaluate the model’s performance and tune its hyperparameters. The test set was used to evaluate the trained model’s performance on unseen data. To reduce the variance of the model’s weights and improve its generalization, the training data was shuffled and divided into small batches of 16 images before each epoch.

The classification CNN was trained on a randomly split dataset consisting of two parts: a training set, and a test set. The training dataset was split into a training and a validation using a ratio of 7:3. The training set was utilized to optimize the model’s parameters, while the validation set was used to evaluate the model’s performance and fine-tune its hyperparameters. The test dataset was used to assess the trained model’s performance on new, unseen data. To improve the model’s generalization and decrease the variance of its weights, the training data was shuffled and split into small batches of 16 images before each epoch. This process was iterated 20 times to ascertain the ideal model and performance tolerances. This comprehensive approach to training and testing the CNN allowed for accurate evaluation of its performance on both seen and unseen data, ensuring the model’s optimized and reliable results. The Adam optimizer (Kingma & Ba, 2014), a stochastic gradient descent method based on adaptive estimation of first and second order moments, was utilized to adjust the model’s parameters in order to minimize the loss function. The categorical cross entropy loss function was chosen for this study because it is often used for classification tasks involving the prediction of discrete class labels. It measures the difference between the predicted probability distribution over the classes and the true distribution and is minimized when the predicted probability for the true class is maximized. The initial learning rate for training was set to 0.001. If the validation loss did not change for 10 epochs, the learning rate was reduced by a factor of 0.05. The models were trained for a total of 400 epochs.

### Model evaluation

Model evaluation is the process of assessing model performance on unseen data, also known as the test set. In this study, the classification model’s performance was evaluated using two evaluation metrics: accuracy and loss. Accuracy is defined as the proportion of correct predictions made by the model, while loss is a measure of the error between the predicted output and the true output. Loss is often used as a training loss to guide the optimization process, and the goal during training is to minimize loss in order to improve the model’s prediction accuracy. The networks were trained and inferred using a notebook that had an Intel Core i9-12900H processor, 64GB of RAM, and an NVIDIA GeForce RTX 3080 Ti graphics card with 16GB of VRAM. Additionally, the code for the workflow is available for public usage on GitHub at: https://github.com/collinswakholi/Potato-defect-detection

This allows for the reproduction of the results obtained in this study and enables other researchers to build upon this work..

## Results

Top-down imaging of whole tubers and flatbed scans of halved tubers was used to collect measurements of tuber size, shape, skin color and flesh color for each clone. Top-down imaging was performed using both a dark non-reflective background and a lightbox background.

### Measurement of tuber size

Systematic differences in the relative size of an object standard were observed across all imaging configurations. Variation in area measurements calculated from a size standard was minimized on the black background: standard deviation for object of same size: 3.97 mm^2^ on the black background, 4.54 mm^2^ on the scanner, and 12.81 mm^2^ on the lightbox background (Fig. S4). Variation in size measurements was generally less than 5% of the total for an object the size of a poker chip (area: 1075 mm^2^, diameter: 37 mm) and linear measurements exhibit a standard error of < 2 mm (Fig. S4). The size standard exhibited significantly less variation on the black background relative to the lightbox in the top-down imaging configuration. Regardless of conformation used, variation in measurement was small relative to the size of the object standard.

Tuber size in terms of length and width as measured by caliper or gravimetric measurement of tuber weight exhibited high correlation with relative pixel area (Fig. 1). Tuber length as measured by caliper and relative pixel area exhibited the highest correlation (r^2^ > 0.98) whereas width measurements (r^2^ > 0.96) and comparison of tuber weight and relative tuber area (r^2^ > 0.93) all exhibited correlation greater than 0.9; indicating that machine vision measurements provide adequate predictive ability to estimate tuber size within this population. Measurements of these size characteristics from different faces/sides of each tuber were practically identical (r^2^ > 0.98; for all characteristics), suggesting that it is not necessary to image both sides of each tuber to measure tuber size traits.

**Fig. 1.**
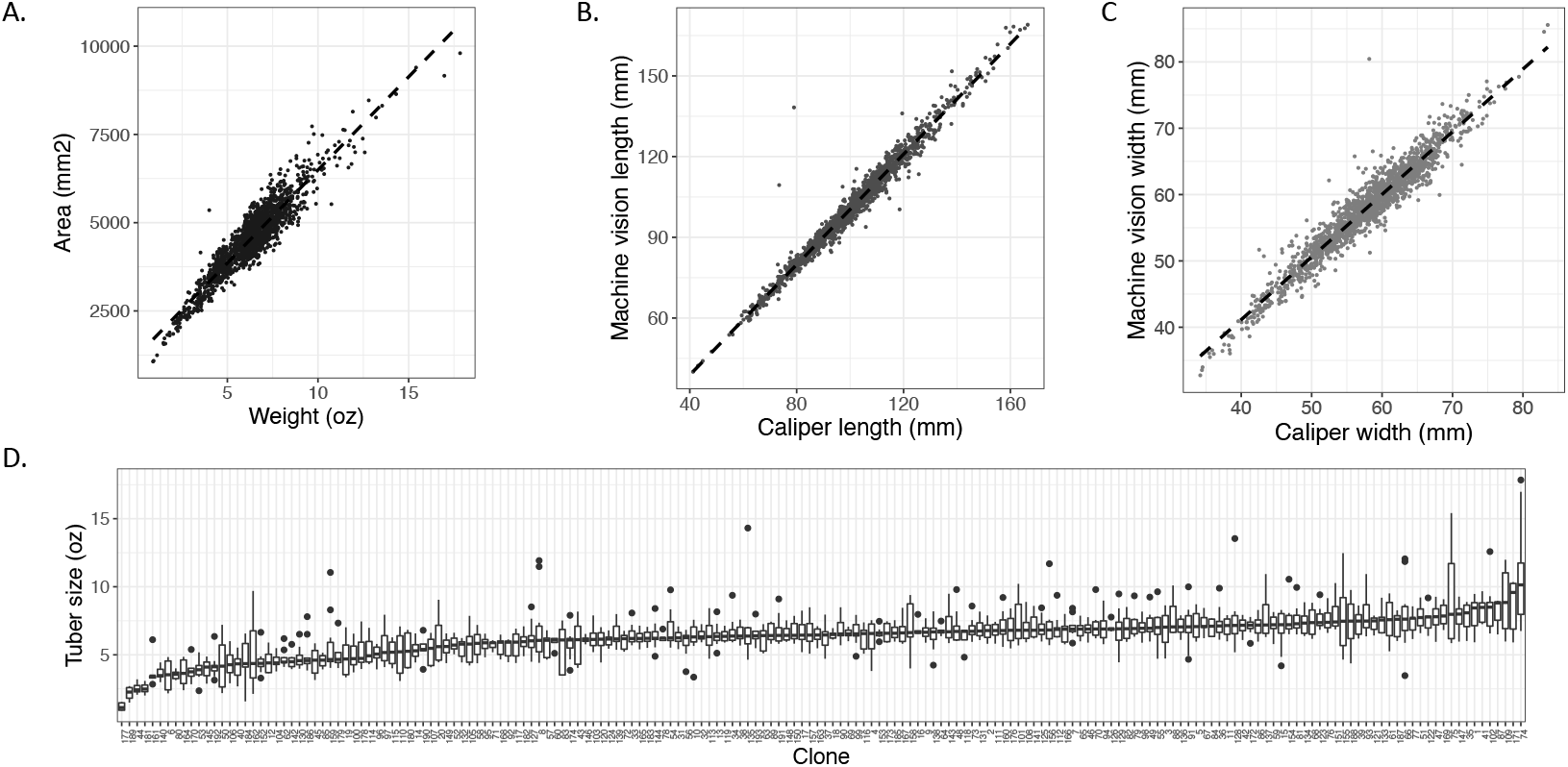
Manual and computer vision measurements of tuber size are highly correlated and exhibit substantial variation within the A08241 population. A) Correlation between tuber weight (oz.) measured gravimetrically and tuber area (mm^2^) as estimated from digital images. B) The relationship between tuber length using calipers (mm) and tuber length as measured using computer vision techniques (mm). C) Correlation between caliper measurements of tuber width (mm) and measurements derived from computer vision (mm). D) The distribution of tuber size across the A08241 population sorted by median of each clone.

**Table 1.**
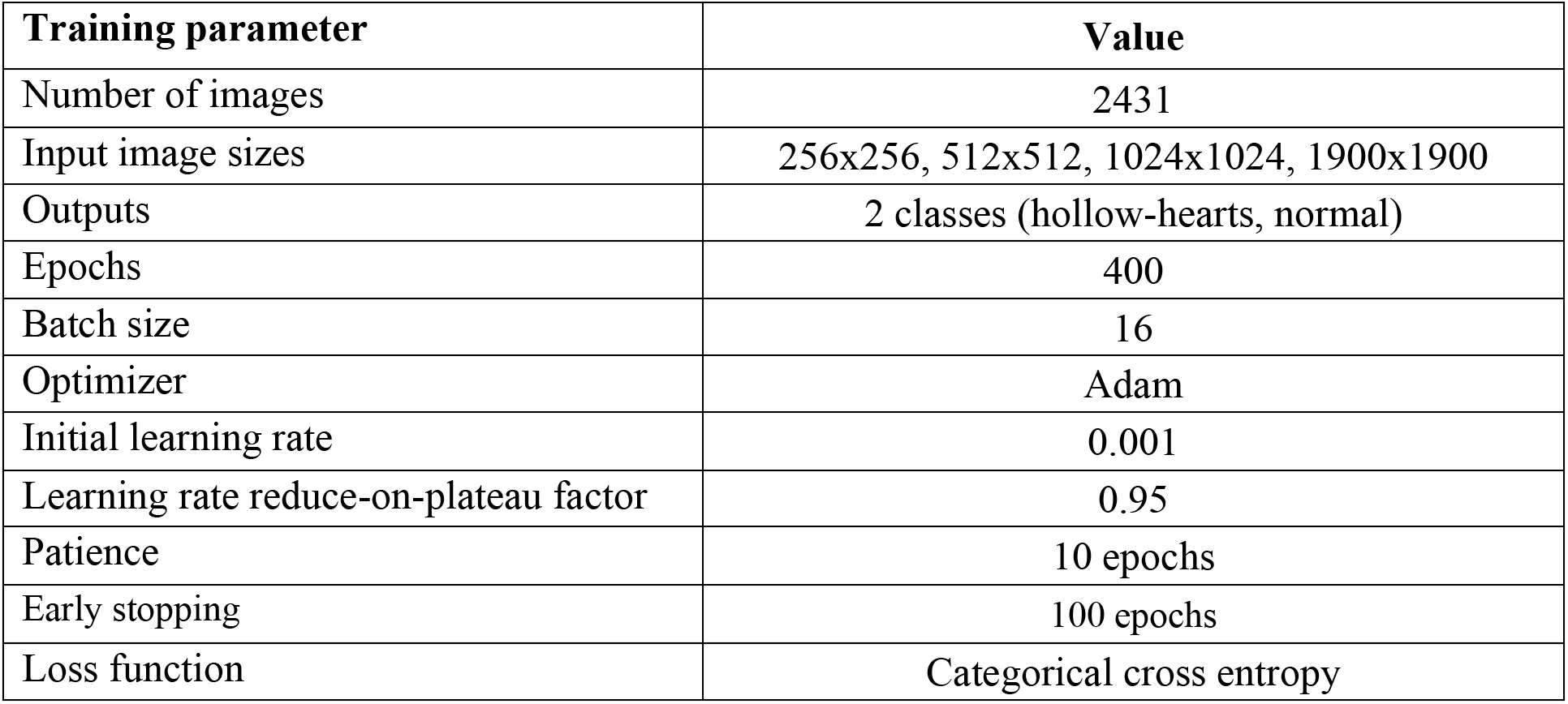
A summary of the model training parameters.

The distribution of mean tuber size (weight and area) from clones in the A08241 population is approximately normal but slightly skewed towards larger tubers (Fig. S5). The mean tuber weight of this population as calculated on a per clone basis is estimated to be 6.33 oz but ranged from 1.19 oz (clone 177) to 10.86 oz (clone 074); whereas the coefficient of variation is ∼16% of the size average across clones (Fig. 1D). Standard deviation was positively correlated with average tuber size (r^2^ = 0.37). All measurements of tuber size exhibited high broad sense heritability (h^2^ > 0.5; Table 2). The heritability of machine vision measurement of tuber characteristics was roughly equivalent to the heritability of manual measurement in every case (Table 2). The standard error of these size measurements decreased as the number of tubers sampled increases but does not exhibit asymptotic behavior at the largest number of tubers sampled (10) suggesting that measuring additional tubers may provide an increasingly accurate estimate of population mean (Fig. S6A).

**Table 2.** Broad-sense heritability of each trait.

| <u>Trait</u> | <u>Broad-sense</u><br><u>heritability</u> | <u>Residual</u><br><u>error</u> |
| --- | --- | --- |
| Tuber weight | 0.54 | 0.46 |
| Tuber area (MV) | 0.55 | 0.45 |
| Tuber length (caliper) | 0.60 | 0.40 |
| Tuber length (MV) | 0.59 | 0.41 |
| Tuber width (caliper) | 0.64 | 0.36 |
| Tuber width (MV) | 0.65 | 0.35 |
| Tuber L/W ratio (caliper) | 0.66 | 0.34 |
| Tuber L/W ratio (MV) | 0.65 | 0.35 |
| SVA | 0.82 | 0.18 |
| Eccentricity | 0.63 | 0.37 |
| PC1.shape | 0.24 | 0.76 |
| PC2.shape | 0.14 | 0.86 |
| PC3.shape | 0.21 | 0.79 |
| PC1.skin | 0.49 | 0.51 |
| PC2.skin | 0.41 | 0.59 |
| PC3.skin | 0.53 | 0.47 |
| PC1.flesh | 0.43 | 0.57 |
| PC2.flesh | 0.53 | 0.47 |
| PC3.flesh | 0.38 | 0.62 |

### Measurement of tuber shape

Variation in the shape of the size standard on the black background was small as measured by both aspect ratio (mean: 1.010, standard deviation: 0.004) and eccentricity (mean: 0.139, standard deviation: 0.029). Both ordinal scoring (aka SolCAP visual assessment, SVA) of tuber shape and caliper measurement of aspect ratio, were used to quantify tuber shape manually, whereas aspect ratio, eccentricity, and principal components values calculated from tuber biomass profiles were obtained using machine vision techniques. Both manual and machine vision measurements of aspect ratio were highly correlated (r^2^ > 0.98) whereas the categorical values of SVA showed substantial correlation (r^2^ = 0.60) but obviously fail to capture the continuous nature of this trait (Fig. 2). The variance explained by principal components (PC) calculated from tuber biomass profiles substantially drops off after the first several PCs (>90% variance explained, Fig. S7). The first PC (PC1_shape_) captures roughly 42.1% of the shape variation observed and is not correlated with aspect ratio (r^2^ = 0.001) or SVA (r^2^ = 0.05) (Fig. 3, Fig. S7). The second PC explains roughly half the variance of PC1_shape_ (PC2_shape_; 22.1% variance explained) and also does not exhibit correlation with aspect ratio (r^2^ = −0.08), whereas PC3_shape_ explains 15.6% of the variation and exhibits a significant correlation with aspect ratio (r^2^ = 0.52) (Fig 3, Fig. S7). Visualizing the variance across these sweeps seems to indicate that PC1_shape_ may describe the variation in tuber thickness at the stem or bud/rose end in this population (Fig. 4a). Examination of tubers exhibiting extreme values along the PC1_shape_ axis supports this conclusion (Fig. 4b). Generally, shape characteristics also exhibited substantial repeatability as measured by correlation between both faces/sides of each tuber. Aspect ratio and eccentricity exhibited the highest correlation between the sides/faces of individual tubers (r^2^ > 0.98), whereas PC scores derived from tuber biomass profiles exhibited slightly lower correlation (PC1_shape_ > 0.93, PC2_shape_ > 0.78, PC3_shape_ > 0.84).

**Fig. 2.**
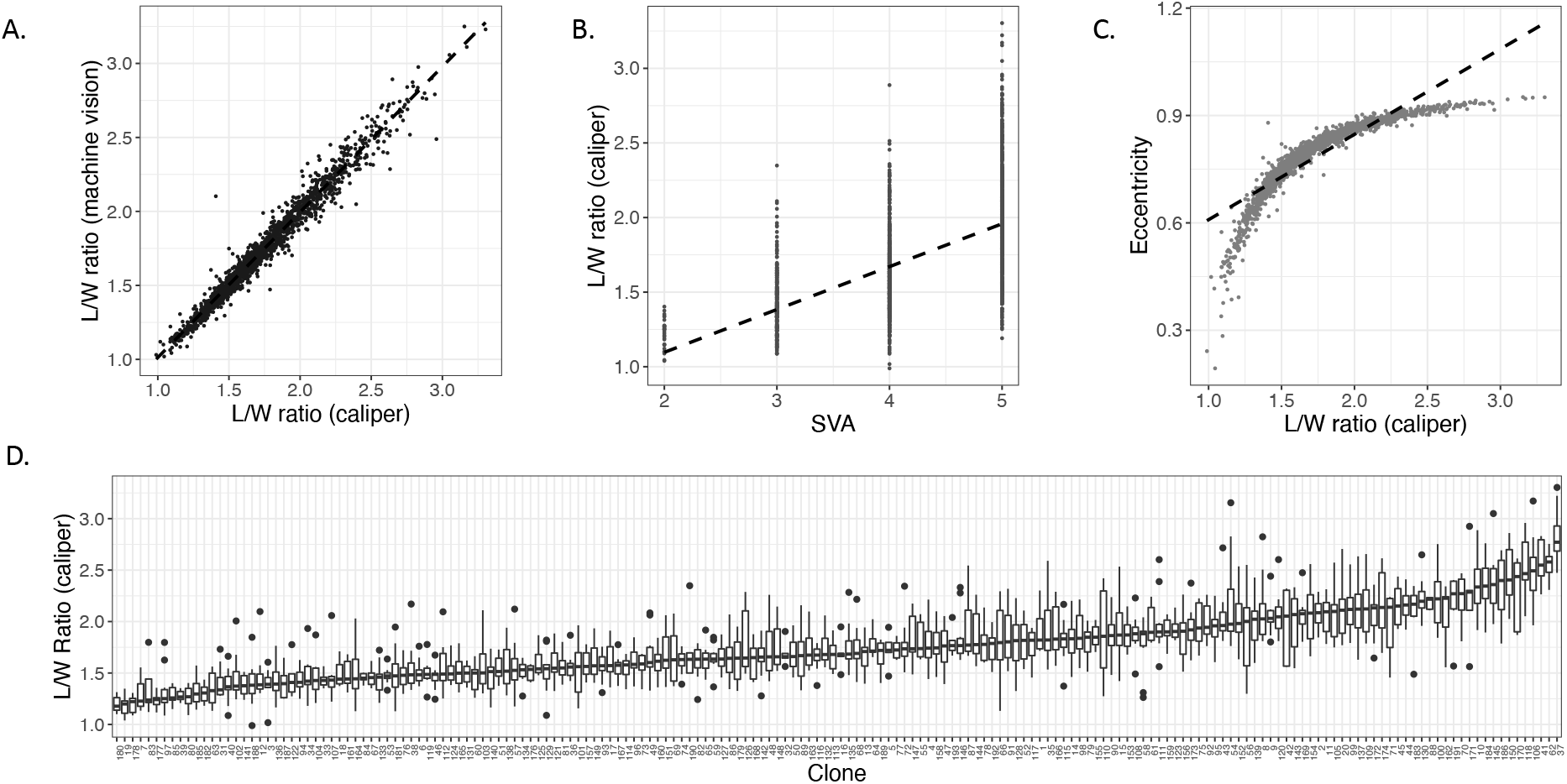
Measurements of tuber shape acquired using manual and computer vision methods are highly correlated and the A08241 population exhibit variation for each of these characteristics. A) Correlation between tuber length/width ratio measured using digital calipers and machine vision estimates extracted from digital images. B) A substantial correlation between length/width ratio and SVA index scoring is observed, but SVA does not accurately reflect the continuous variation in this trait. C) Eccentricity of the bounding ellipse as measured by computer vision measurements is correlated with length/width ratio but the relationship between these measurements isn’t as strong at the tails of the distribution. D) The distribution of aspect ratio within the A08241 population sorted by the median of each clone.

**Fig. 3.**
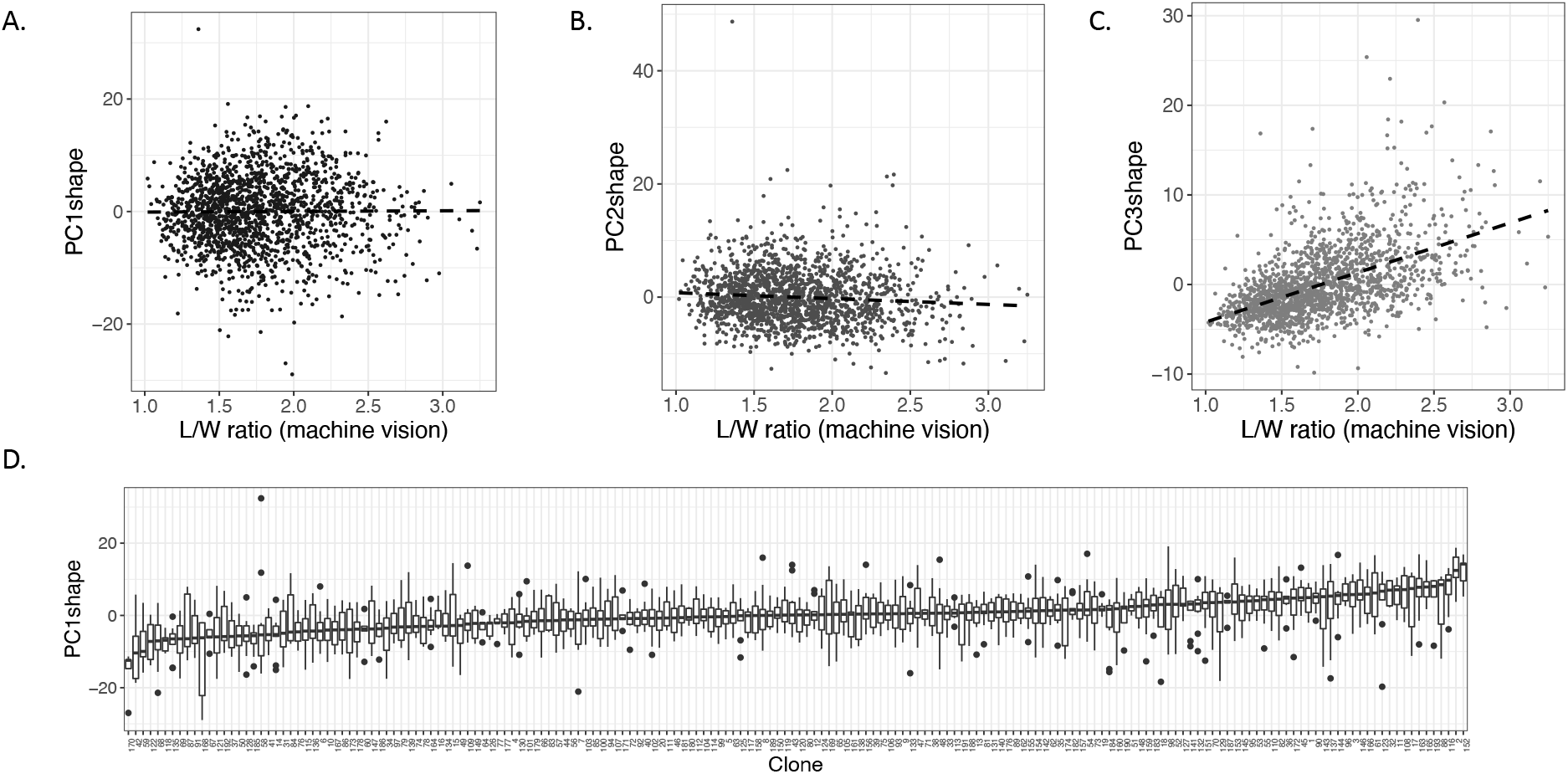
Principal component 1 and principal component 2 derived from tuber biomass profiles (PC1_shape_) are not correlated with length/width ratio and exhibits variation in the A08241 population. A) PC1_shape_ is not correlated with tuber length/width ratio. B) PC2_shape_ exhibits a no correlation with tuber length/width ratio. C) PC3_shape_ exhibits a positive correlation with length/width ratio. D) The distribution of PC1_shape_ within the A08241 population sorted by the median of each clone.

**Fig. 4.**
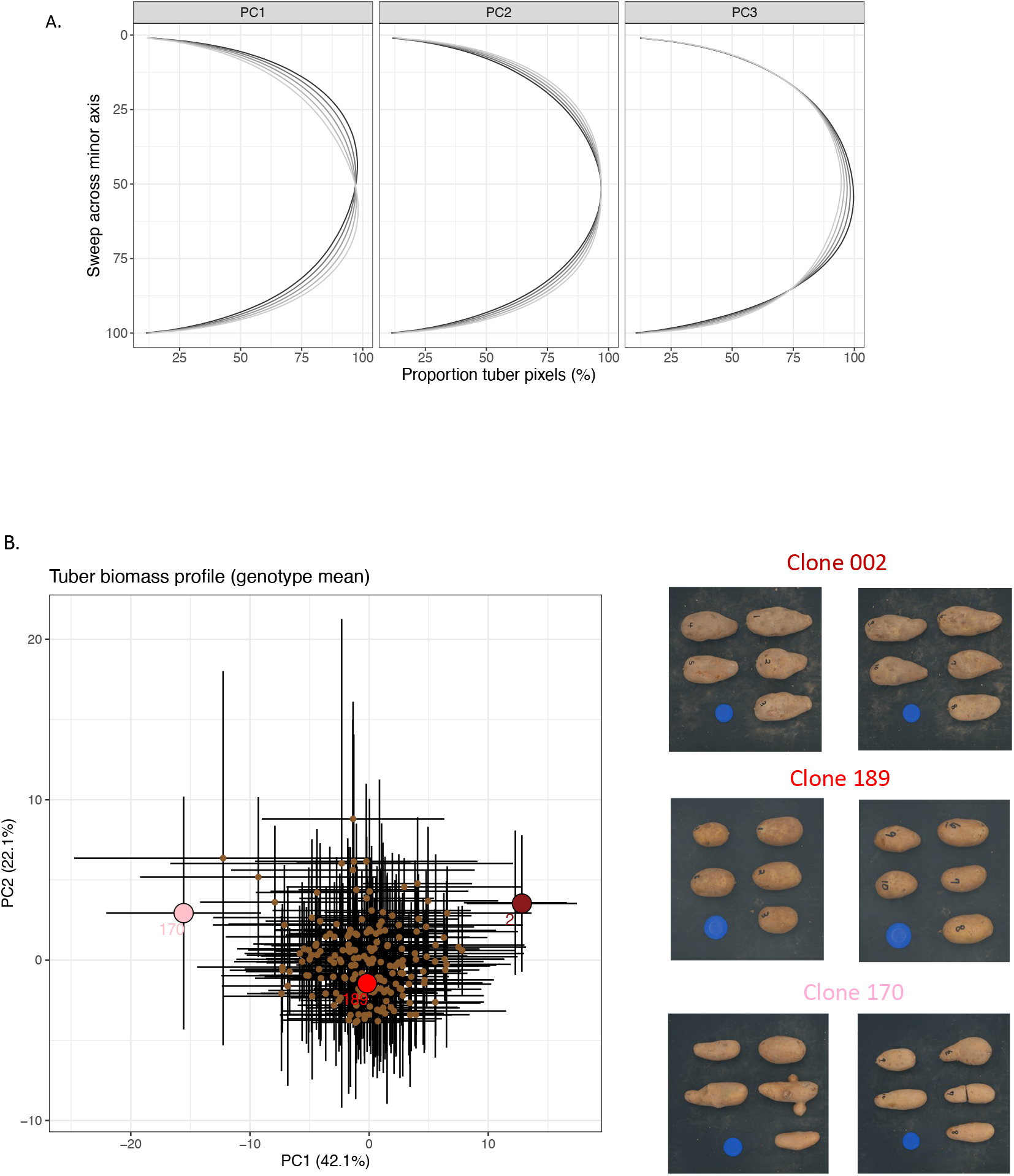
Dimensional reduction of tuber biomass profiles using principal component analysis suggests that these latent traits can provide quantitative information about tuber shape not captured by aspect ratio. A) Eigenvectors for principal components analysis of potato tuber shape after normalization for aspect ratio. Lines represent parameter sweep of three principal components, capturing symmetrical variation in tuber shape. B) The values of principal component 1 plotted against the values of principal component 2 (clone average), where the tubers exhibiting extreme values (max, mean, min) are plotted in different shades of red corresponding to their image label.

The distribution of mean aspect ratio from clones in the A08241 population is unimodal but likely non-normal and skewed towards larger tubers (Fig. S5). The mean tuber aspect ratio within this population as calculated on a per clone basis is 1.76 but ranged from 1.17 (clone 019) to 2.81 (clone 037); whereas the standard deviation around the mean within each clone averages 0.21 across the population. Aspect ratio exhibits substantial heteroskedasticity, as the correlation between trait value and standard deviation is substantially correlated (r^2^ > 0.57).

Broad sense heritability was greatest for SVA (H^2^ = 0.82), and the heritability of aspect ratio as measured manually (H^2^ = 0.66) and using machine vision (H^2^ = 0.65) were almost identical (Table 2). Heritability of tuber biomass profile PCs suggest that the first three PCs may be influenced by genetics (all have H^2^ > 0.14) but little evidence exists for a genetic signal beyond PC4 (all have H^2^ < 0.11).

Substantial differences in variation were observed between traits. SVA exhibited the smallest standard error, whereas aspect ratio and eccentricity expressing a medial value, and PC_shape_ values derived from tuber biomass profiles displayed the largest standard error (Fig. S6b). As observed for the tuber size measurements, the standard error of each measurement type decreased as the number of tubers sampled increased. The rate at which the standard error of SVA, aspect ratio, and eccentricity decrease with additional tubers sampled appears to approach an asymptote after 8 tubers are sampled, suggesting that measurement of additional tubers will provide diminishing returns.

### Measurement of tuber skin and flesh color

Colorimetric values derived from a color checker standard (Fig. S8) were compared within and between images acquired using a flatbed scanner (Fig. S9), top-down imaging on the black background (Fig. S10), and top-down imaging on a light box (Fig. S11). Principal component analysis (PCA) was used to perform dimensional reduction across the mean value of each RGB channel and the values of each color chip were compared. The standard deviation (SD) of colorimetric values derived from the color checker along PC1 was almost three times greater on the black background (SD = 0.56), and more than three and a half times larger on the light box (SD = 0.66) relative to the values extracted from the flatbed scanner (SD = 0.19). Color correction of the top-down images collected on the black background and light box reduced the standard deviation substantially (SD = 0.08 and SD = 0.15; respectively). Across all three image acquisition backgrounds, standard deviation was generally largest in magnitude among lighter colors (colors with higher values on each of the R, G, B channels) relative to darker colors (lower RGB values). Along PC1, derived from non-color corrected images, color checker chips 19, 5, 4, 23, and 6 were the most variable; whereas variation was minimized for chips 1, 15, 24, 12, and 2 (Fig. S8, Table S1). Color correction had a substantial influence on this variation. Within color corrected images chips 6, 1, 9, 20, and 19 exhibited the largest variation along PC1, whereas chips 21, 10, 17, 13, and 24 exhibited the least variation (Fig. S8, Table S1)

Color corrected, top-down images of intact tubers placed on a black background and uncorrected images of halved tubers collected using the flatbed scanner were used to assess tuber skin and tuber flesh characteristics. For both tuber skin and tuber flesh, the mean signal value of each tuber contour within the R, G, and B color channels was tabulated and PCA was used to reduce the dimensionality of the data. The proportion of variance captured by PCA decreased rapidly for both tuber skin and flesh color (Fig. S6) with the first PC capturing greater than 87% of the variation in both datasets. Repeatability of skin color, measured as the correlation between values calculated from individual sides/faces of each tuber, was moderate-to-high (PC1_skin_ = 0.71, PC2_skin_ = 0.64, PC3_skin_ = 0.82). Repeatability of flesh color measurements could not be calculated, as only a single half of each tuber was imaged.

PC1 derived from colorimetric data likely captures the reflective features of both skin and flesh color (Fig. 5). Values of PC1_skin_ and PC1_flesh_ were most highly correlated with L* (lightness) channel from the CIE LAB color space (Table S2). Coloration of tuber skin along PC1 (PC1_skin_) exhibited a unimodal distribution centered around zero (Fig. S5). Clone 176, 170, 172, and 20 can be found at the far right of the PC1_skin_ distribution and had the lightest skin (average values greater than 2.5), whereas clones 088, 058, and 083 exhibited the darkest tuber skin with average PC1_skin_ values less than −2.5 (Fig. 5). PC1 derived from colorimetric values of tuber flesh (PC1_flesh_) exhibited a similar distribution, mean, and color trends as was observed for tuber skin. Tubers with the lightest flesh: clones 070, 141, 001 and 022, exhibit average values of PC1_flesh_ greater than 3.3; whereas clones 193, 097, 100, and 078, exhibit the darkest flesh in this population with PC1_flesh_ values less than −2.3 (Fig 5). No clear trends discernable by the human eye were observed along PC2, PC3, etc. for either tuber skin or flesh color.

**Fig. 5.**
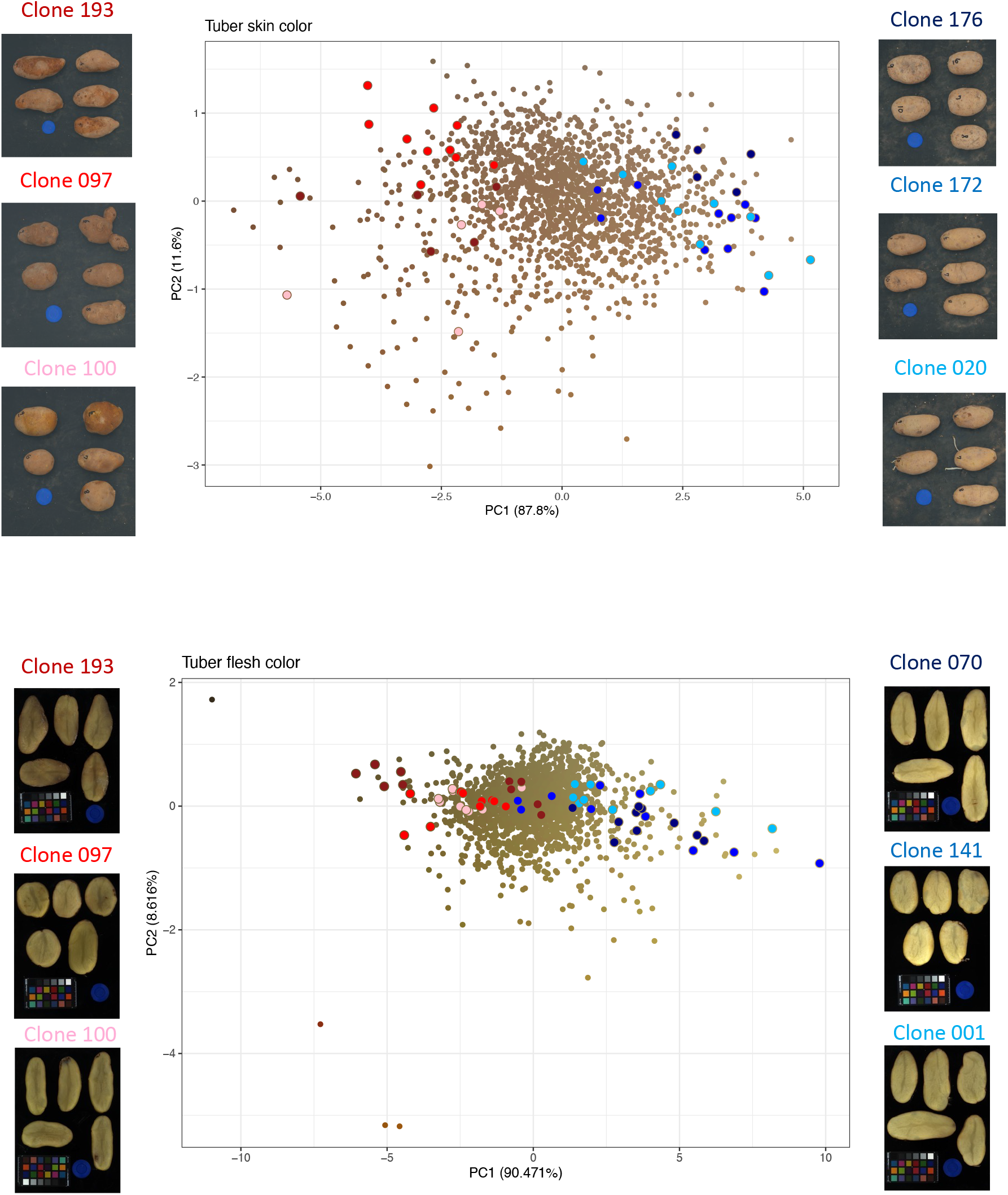
Dimensional reduction of colorimetric values performed using principal component analysis suggests that the major source of phenotypic variation in tuber skin and tuber flesh relates to the light reflective properties of color in both tissue types. A) PC1_skin_ exhibits a spectrum of color values that progresses from dark to light along the x-axis. B) The color values of tuber flesh along PC1_flesh_ exhibit a trend of progression from dark to light along the x-axis.

Broad-sense heritability along both PC1_skin_ and PC1_flesh_ measurements was high. PC1_skin_ measurement of tuber color in this population exhibited a broad-sense heritability of 0.49, whereas PC2_skin_ and PC3_skin_ exhibited H^2^ values of 0.41 and 0.53 respectively. PC1_flesh_ exhibited a broad-sense H^2^ value of 0.43. This value was lower than the H^2^ value observed for PC2_flesh_ (0.53) but higher than the value observed for PC3_flesh_ (0.38).

Outside of PC3_skin_, the standard error of the colorimetric measurements decreased as the number of tubers sampled increases (Fig. S6C). The first PC from both skin and flesh tissue captures a majority of the variation in this colorimetric data and also expresses the largest standard error. The reduction of standard error in PC1_flesh_, PC2_flesh_, PC3_flesh_, and PC1_skin_ is not clearly approaching an asymptote, suggesting that measuring additional tuber will provide a better estimate of sample mean. PC2_skin_ and PC3_skin_ on the other hand, exhibit asymptotic behavior after four tubers have been sampled.

### Classification of hollow heart defect

A total of 50 individual tubers from 27 clones were identified to exhibit hollow heart defect. The frequency of this defect among affected clones varied with slightly more than half of the affected clones exhibiting a single defective tuber (55%) and a single clone exhibiting 8 hollow heart affected tubers out of 10 total. The CNN model was trained using the parameters in Table 1 for 400 epochs. Model accuracy improves and loss decreases with increasing training epochs, indicating that the model is learning and becoming more effective at the task (Fig. 6). However, after around 60 epochs, the trend of both the accuracy and loss appear to plateau, meaning that the model’s performance stops improving significantly with more training (Fig. 6). This plateau may occur because the model has reached a local minimum in the loss function and is no longer able to make significant progress on the task, or it may be due to overfitting, where the model is too closely tuned to the specific training data and is not able to generalize well to new, unseen data. This same trend was observed while training the model on all the different resolution datasets. We evaluated the performance of deep learning models on images of different resolutions, with the goal of assessing the generalizability of the models. The model performance in terms of accuracy, loss, confidence score, and inference time are presented in Table 3, and Fig. S12. The results show that training the models on images with a resolution of 512×512 pixels resulted in the highest accuracy (an average of 99.12% and a maximum of 100%), lowest loss (an average of 0.0366 and a minimum of 0.0027), and highest confidence score (an average of 99.65% and a maximum of 100%) when compared to other resolutions. Examples of accurate predictions and common misclassifications observed in models with less than 100% accuracy can be observed in Fig. 7 and Fig. S13 respectively. While the 256×256 resolution performed well in terms of precision (see Table 3, Fig. S12), it failed to capture the level of detail necessary for accurate classification. In contrast, using higher resolutions such as 1024×1024 and 1900×1900 pixels did not improve the model performance, possibly due to limitations in the model architecture that made it unsuitable for handling larger images. These findings suggest that there is a trade-off between image resolution and model performance, and choosing the appropriate resolution is crucial for achieving optimal results. In summary, our study highlights the importance of carefully considering the image resolution when training deep learning models to ensure their generalizability and performance on a variety of image types.

**Fig. 6.**
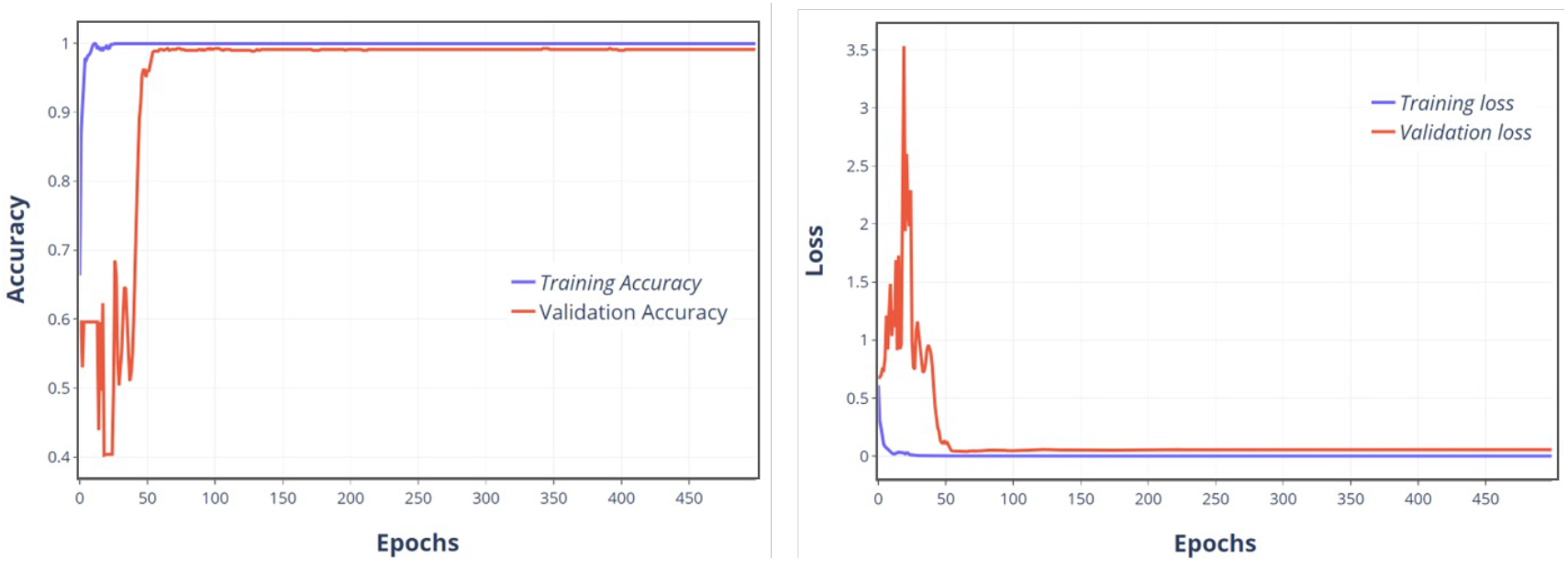
Trends of accuracy and loss during model training. The x-axis represents the training epochs, and the y-axis displays the accuracy or loss. The training results are plotted in blue, while validation results are plotted in orange.

**Fig. 7.**
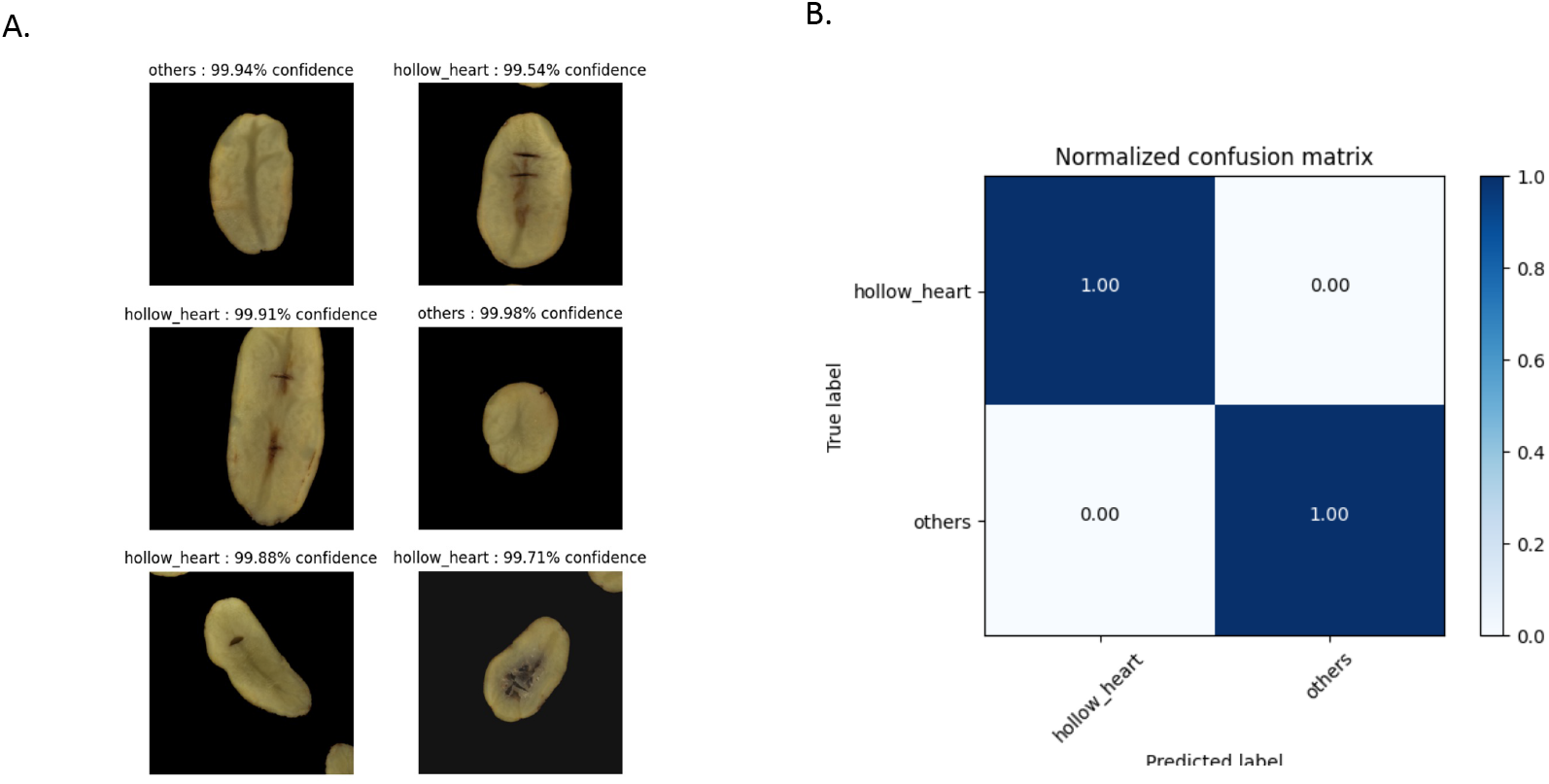
CNN model predictions on test dataset. A) Examples of subjects classified as having hollow heart and those falling in category ‘other’. B) Normalized confusion matrix of results from the best performing model.

**Table 3.** Model performance on images with different resolution.

| Resolution | # of Models | Test Accuracy (%) |  |  | Test Loss |  |  | Model Confidence (%) |  |  | Inference Time (ms) |
| --- | --- | --- | --- | --- | --- | --- | --- | --- | --- | --- | --- |
|  |  | Range | Mean | Std. | Range | Mean | Std. | Range | Mean | Std. |  |
| $256 \times 256$ | 20 | 99.57-97.45 | 98.97 | 0.485 | 0.0915-0.0228 | 0.0422 | 0.01594 | 100.00-50.00 | 99.53 | 2.772 | 2.33<br>0.33 |
| $512 \times 512$ | 20 | 100.00-95.74 | 99.12 | 0.934 | 0.2289-0.0027 | 0.0366 | 0.04723 | 100.00-50.00 | 99.65 | 2.178 | 2.94<br>0.16 |
| $1024 \times 1024$ | 14 | 100.0-94.26 | 98.66 | 1.576 | 0.1613-0.0016 | 0.0396 | 0.04147 | 100.00-50.00 | 99.43 | 2.988 | 7.32<br>0.28 |
| $1900 \times 1900$ | 12 | 99.57-92.13 | 96.33 | 2.292 | 0.2182-0.0203 | 0.0971 | 0.05566 | 100.00-50.00 | 97.28 | 8.372 | 21.30<br>1.09 |

## Discussion

This study provides further support to the compendium of evidence suggesting machine vision is a reliable and effective tool to measure many independent phenotypic characteristics of potato tubers (Caraza-Harter & Endelman, 2020; Liu et al., 2021; Miller et al., 2023; Neilson et al., 2021; Rady & Guyer, 2015; Si et al., 2017; Su et al., 2017). To our knowledge, this is the first study that has applied machine vision techniques to estimate tuber size, shape, external and internal colorimetric characteristics as well as classify defects in a replicated, medium-to-large size biparental potato linkage mapping population. Our results indicate that measurement precision of a size standard is maximized when images can be acquired on a non-reflective background, but that choice of background doesn’t contribute as a major source of variation so long as tubers exhibit adequate contrast relative to the background for the purpose of threshold isolation. Estimation of colorimetric values from a color checker standard demonstrates that substantial measurement variation associated with color quantification should be anticipated; and that lighter, more reflective colors will exhibit greater measurement variation relative to darker, less reflective colors.

As reported by several other researcher groups (Hasankhani & Navid, 2012; Liu et al., 2021; Miller et al., 2023; Neilson et al., 2021; Noordam et al., 2000; Si et al., 2017; Su et al., 2017; Zhou et al., 1998), machine vision measurement of size and shape are highly correlated with manual gravimetric, caliper measurement, or index scoring of these characteristics (all characteristics exhibit r^2^ > 0.9). Although a high correlation between relative tuber area (estimated based upon calibration using a size standard) and gravimetric measurements of tuber size was observed in the A08241 population, this trend may not be as robust in other populations where tuber size exhibits greater variation along the z-axis, which cannot be quantified from the 2-dimensional images captured in this study. In the future, these platforms could potentially be improved by incorporating a repeating calibration pattern to improve measurement accuracy (Tabb et al., 2020) and multiple cameras (Tabb et al., 2019) or depth sensors to estimate object length along the z-axis (Liu et al., 2021; Su et al., 2017).

Benefits of machine vision measurement are particularly apparent with regards to quantification of tuber shape and colorimetric features. Results from this study suggest that latent traits, like PCs derived from tuber biomass profiles, may capture additional heritable shape characteristics that are independent (uncorrelated with) of aspect ratio typically quantified by ordinal scoring or manual measurement. Colorimetric features are notoriously difficult to quantify and are often described categorically. Although categorical description provides actionable information, this type of ordinal data may not provide adequate resolution for genetic mapping in linkage and association mapping populations like the A08241 population. An additional consideration is that principal component data derived from shape or colorimetric features is experiment specific, meaning that results and conclusions from this study may not be directly relatable to studies performed on other populations unless data are analyzed together as a single dataset.

Images are a rich source of quantitative data that can be acquired by non-experts using inexpensive, consumer grade sensors. The total cost to construct the system described is less than $2,000 U.S. dollars and the image data collected for this study was collected by an undergraduate student. Staging of tuber samples is fairly labor intensive so delivery of tubers to the sensor using a conveyor system may greatly enhance throughput. The major benefits of collecting this data type are that researchers can extract and unbiasedly quantify many tuber characteristics simultaneously (far more than could be collected by a human in the same timespan) and the data is long lived, meaning that it can always be revisited later to perform additional analysis (extract new features) after the tubers have been discarded. Features including tuber blemishes, physiological defects, or pathogen disease state can be annotated within image datasets like this one and ultimately used to build classification models to predict the presence of defects with minimal human supervision (Liu & Wang, 2021; Marino et al., 2019; Miller et al., 2023; Moallem et al., 2013; W. Zhang et al., 2019). Our study suggests that CNN machine learning techniques are highly effective at classifying the hollow heart defect of potato. The performance of this model was consistent with that of similar studies (Huertas-Tato et al., 2022; Lu et al., 2021; Marino et al., 2019; Moallem et al., 2013; Wang & Xiao, 2021). However, it is worth noting that the model’s performance did not improve with an increase in image resolution used for training. This suggests that the model may benefit from additional training data, better architecture, or regularization techniques to improve its generalization performance and reduce overfitting, especially for larger image sizes. Overall, the classification CNN developed in this study demonstrates great potential as a tool for identifying and classifying potatoes with hollow-heart. The resulting image data can also be used to calculate additional latent trait values (combinations of high-dimensional features that define a group of individuals) to better understand how genotypes differ or respond to environmental perturbation (Feldmann et al., 2020; Gage et al., 2019; Ubbens et al., 2020).

Trait value distribution and estimates of broad-sense heritability suggest that tuber size, length-width ratio, tuber end thickness (PC1_shape_), tuber skin (PC1_skin_) and internal colorimetric features (PC1_flesh_) are highly heritable and are likely inherited as polygenic traits. Use of machine learning models indicates that identification of tuber defects is also possible given a large and diverse enough training set to build accurate classification models. Application of these methods, combined with modern genotyping techniques and quantitative genetics methods will certainly improve our understanding of the genetic features underlying these traits and may help us better predict the characteristics of offspring in potato breeding populations.

## Conclusions

Machine vision measurement of tuber characteristics is an inexpensive, high-throughput method to accurately and unbiasedly quantify size, shape, colorimetric features and quality defects simultaneously. Heritable variation of these characteristics is observed within the A08241 population and the values reported can be directly used for genetic mapping once marker information becomes available for the individual clones. The workflows used to collect this data are well-suited for characterization of early generation breeding material (linkage and association mapping populations) and can be deployed in breeding programs of any size with minimal start up and operating costs.

## Supporting information

AllFigures

AllTables

## Data availability

The image datasets collected during this study are available to the public through the USDA National Agricultural Library Ag Data Commons.

## Acknowledgements

The authors thank Brian Schneider, Darren Hall, Mark Fristad, Ian Fullmer, Jericho Watson, and Russell Elswick USDA-ARS, Aberdeen, ID for their contributions to the collection of data on population A08241 at the Idaho trial site.

## Supplemental Tables

**Table S1.**
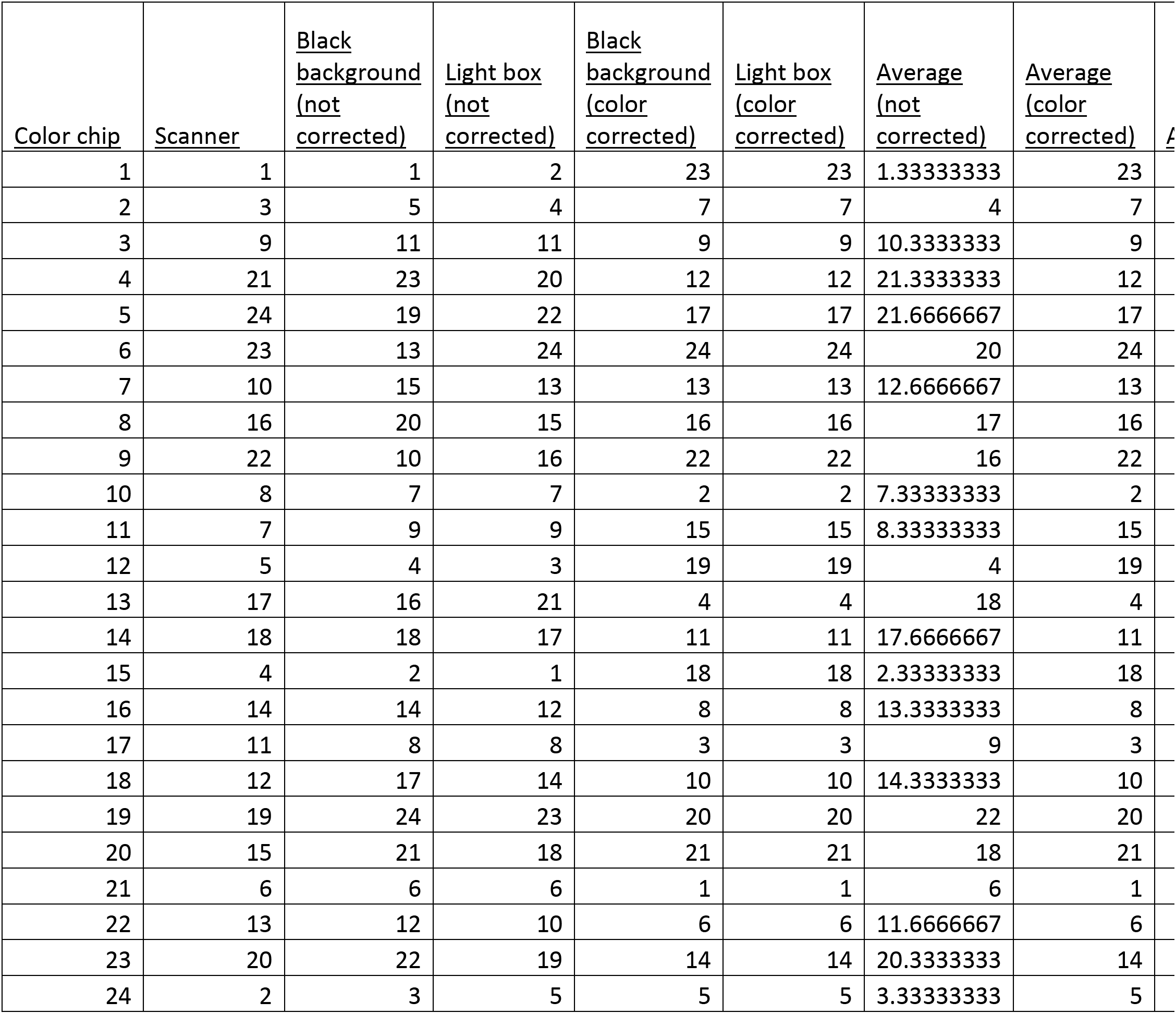
The rank (lowest =1 to highest =24) of standard deviation of reflectance calculated from each color chip on the flatbed scanner, non-reflective black background, and lightbox both with and without color correction.

| <u>Color chip</u> | <u>Scanner</u> | <u>Black background (not corrected)</u> | <u>Light box (not corrected)</u> | <u>Black background (color corrected)</u> | <u>Light box (color corrected)</u> | <u>Average (not corrected)</u> | <u>Average (color corrected)</u> | <u>A</u> |
| --- | --- | --- | --- | --- | --- | --- | --- | --- |
| 1 | 1 | 1 | 2 | 23 | 23 | 1.33333333 | 23 |  |
| 2 | 3 | 5 | 4 | 7 | 7 | 4 | 7 |  |
| 3 | 9 | 11 | 11 | 9 | 9 | 10.33333333 | 9 |  |
| 4 | 21 | 23 | 20 | 12 | 12 | 21.33333333 | 12 |  |
| 5 | 24 | 19 | 22 | 17 | 17 | 21.66666667 | 17 |  |
| 6 | 23 | 13 | 24 | 24 | 24 | 20 | 24 |  |
| 7 | 10 | 15 | 13 | 13 | 13 | 12.66666667 | 13 |  |
| 8 | 16 | 20 | 15 | 16 | 16 | 17 | 16 |  |
| 9 | 22 | 10 | 16 | 22 | 22 | 16 | 22 |  |
| 10 | 8 | 7 | 7 | 2 | 2 | 7.33333333 | 2 |  |
| 11 | 7 | 9 | 9 | 15 | 15 | 8.33333333 | 15 |  |
| 12 | 5 | 4 | 3 | 19 | 19 | 4 | 19 |  |
| 13 | 17 | 16 | 21 | 4 | 4 | 18 | 4 |  |
| 14 | 18 | 18 | 17 | 11 | 11 | 17.66666667 | 11 |  |
| 15 | 4 | 2 | 1 | 18 | 18 | 2.33333333 | 18 |  |
| 16 | 14 | 14 | 12 | 8 | 8 | 13.33333333 | 8 |  |
| 17 | 11 | 8 | 8 | 3 | 3 | 9 | 3 |  |
| 18 | 12 | 17 | 14 | 10 | 10 | 14.33333333 | 10 |  |
| 19 | 19 | 24 | 23 | 20 | 20 | 22 | 20 |  |
| 20 | 15 | 21 | 18 | 21 | 21 | 18 | 21 |  |
| 21 | 6 | 6 | 6 | 1 | 1 | 6 | 1 |  |
| 22 | 13 | 12 | 10 | 6 | 6 | 11.66666667 | 6 |  |
| 23 | 20 | 22 | 19 | 14 | 14 | 20.33333333 | 14 |  |
| 24 | 2 | 3 | 5 | 5 | 5 | 3.33333333 | 5 |  |

**Table S2.**
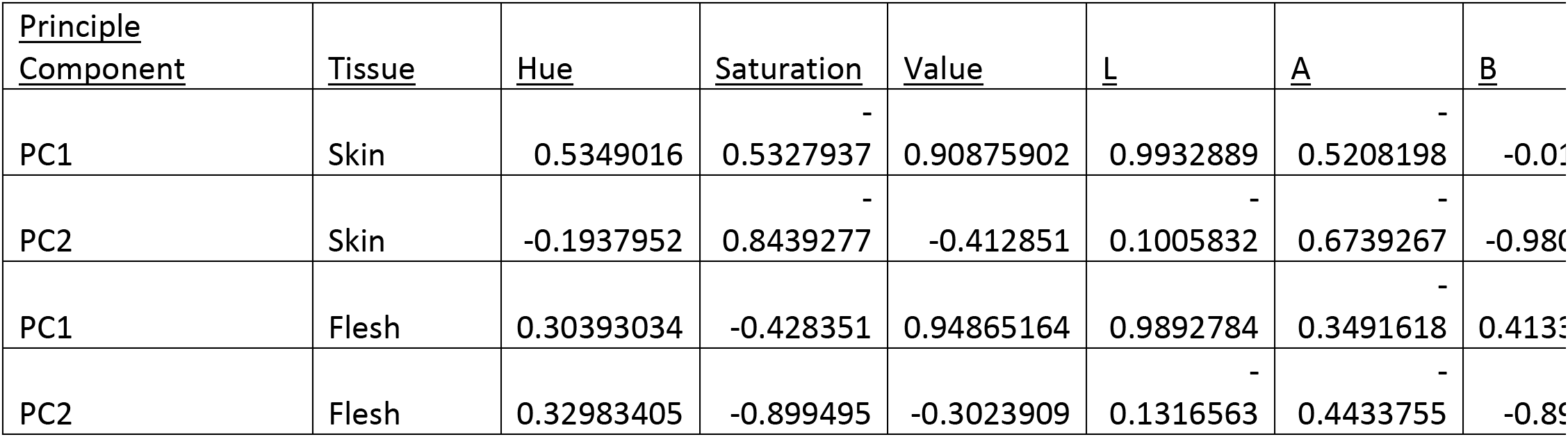
Correlation between principal component values calculated from potato tuber skin and flesh and mean values from HSV and CIE LAB color channels.

## Supplemental Figures

**Fig. S1.**
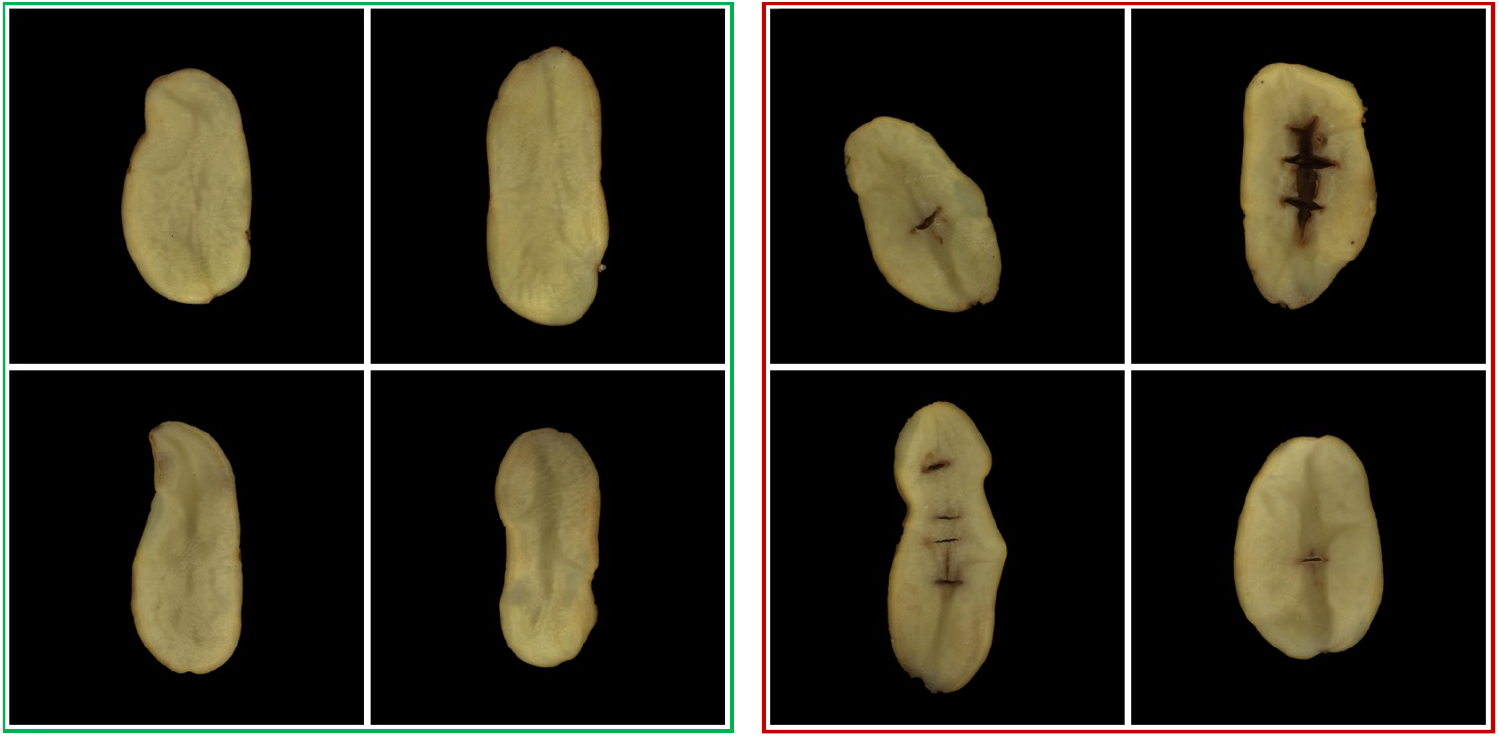
Sample images from the dataset, including a hollow-heart defect potato slice on the right and a normal potato slice on the left.

**Fig. S2.**
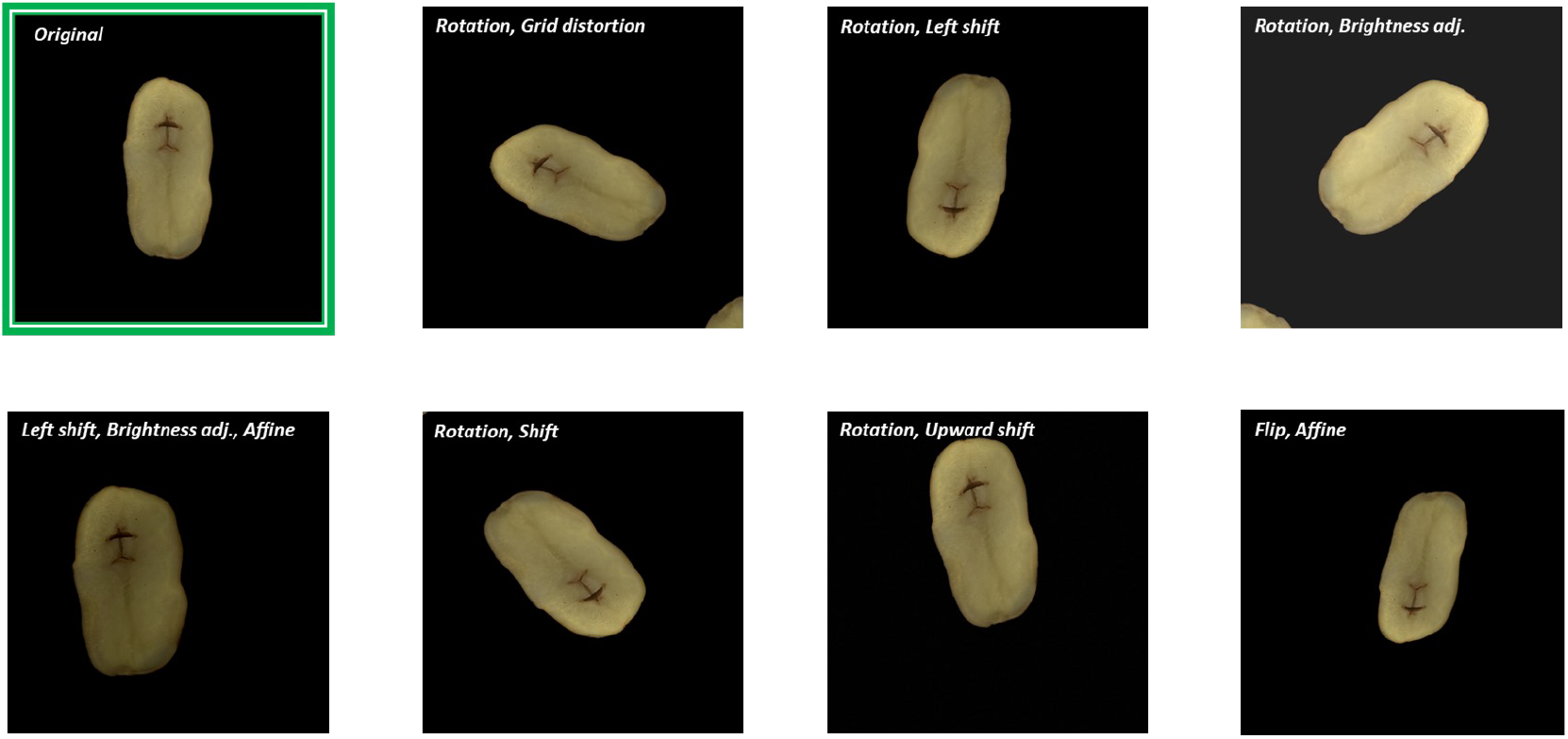
Example of Hollow-heart defect potato slices with image augmentation applied.

**Fig. S3.**
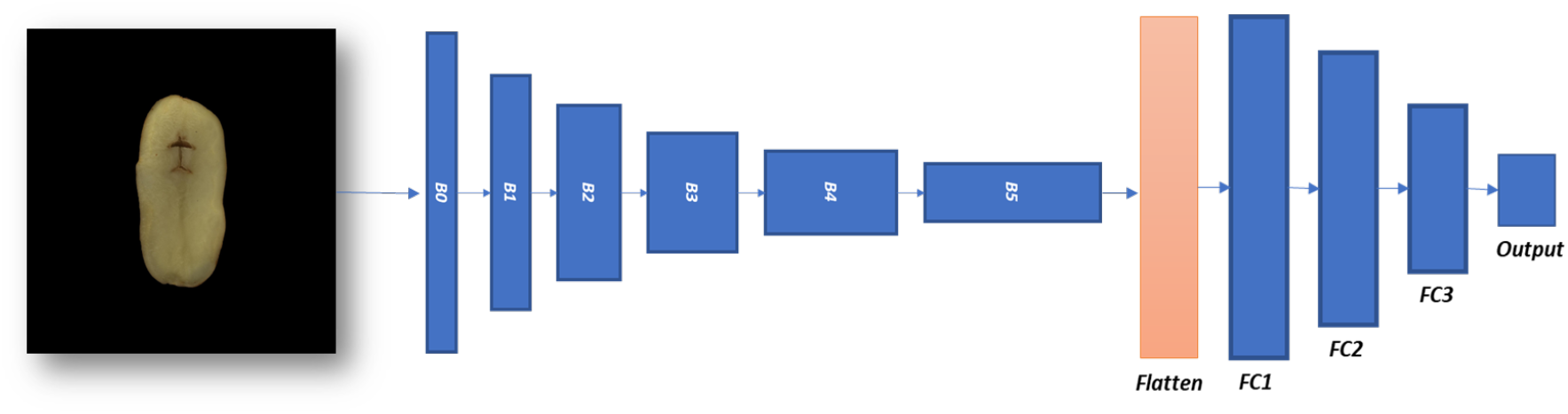
Image classification CNN Architecture used for the classification of hollow-heart defects and normal potatoes.

**Fig. S4.**
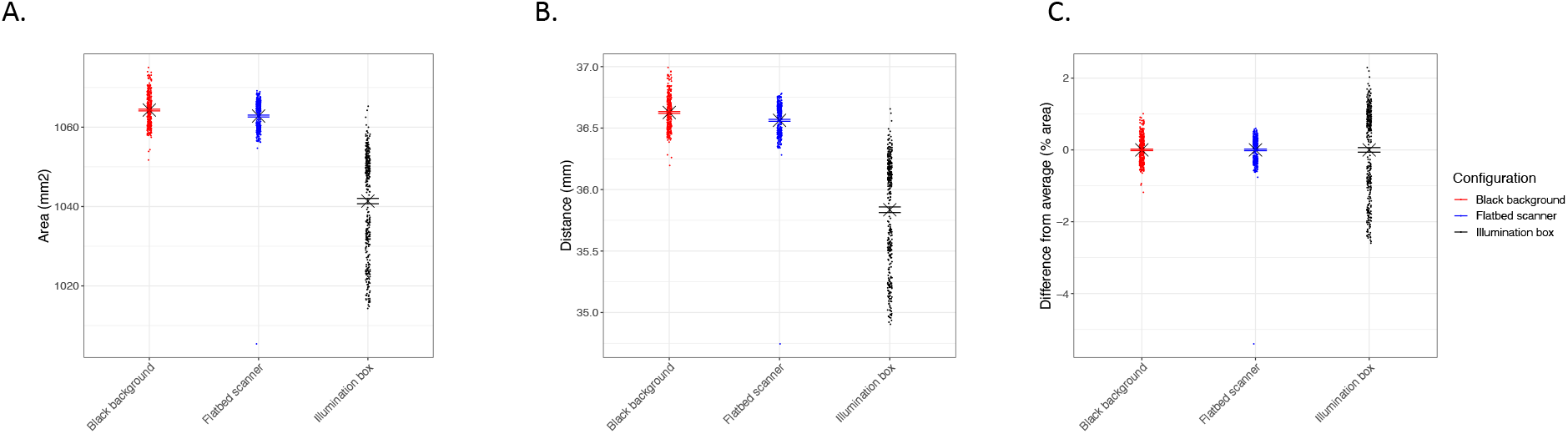
Measurements of a size standard across three different imaging configurations suggest significant differences between configurations exist but all exhibit negligible error. A) Significant differences in size (pixel area) associated with imaging configuration are observed and measurement error is minimized on the flatbed scanner and non-reflective black background. B) Distance measurements exhibited significant differences across all three imaging platforms. Error was minimized on the flatbed scanner. C) Proportionally the % of error observed in measurement of the size standard was greatest when objects were imaged on the illumination box, however all imaging configurations generally exhibited less than 5% error around the true values of the size standard.

**Fig. S5.**
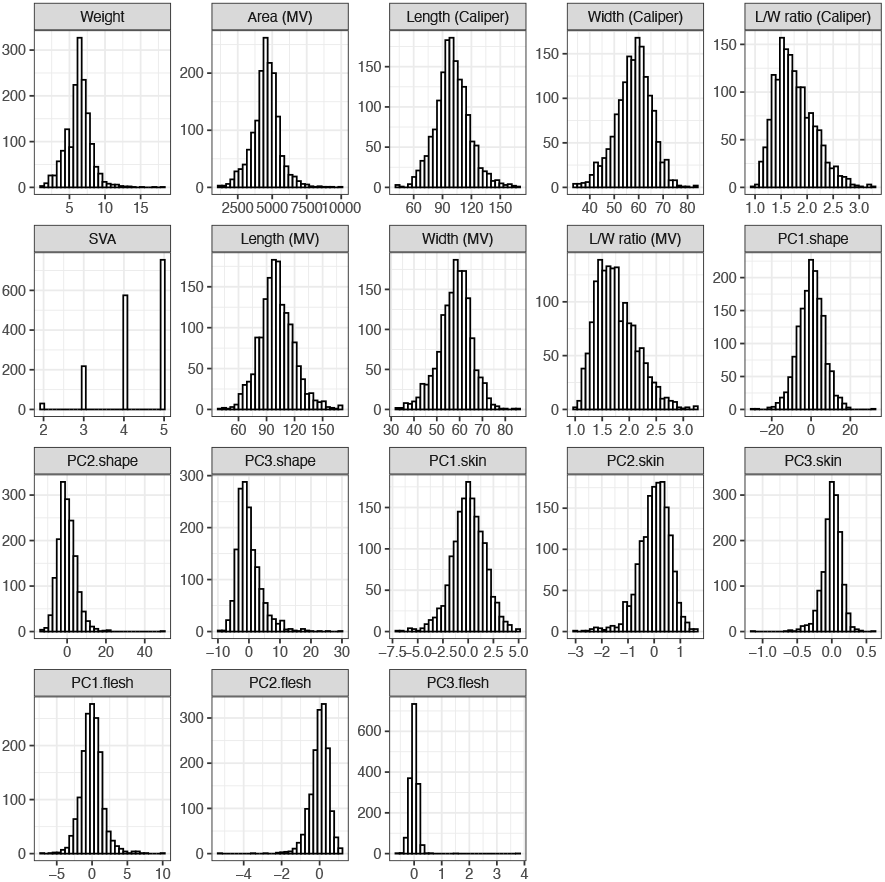
Histograms of the distribution of all traits presented during this study [Digital caliper measurement (Caliper), Machine vision measurement (MV), SolCAP Visual Assessment (SVA), Principal component (PC)].

**Fig. S6.**
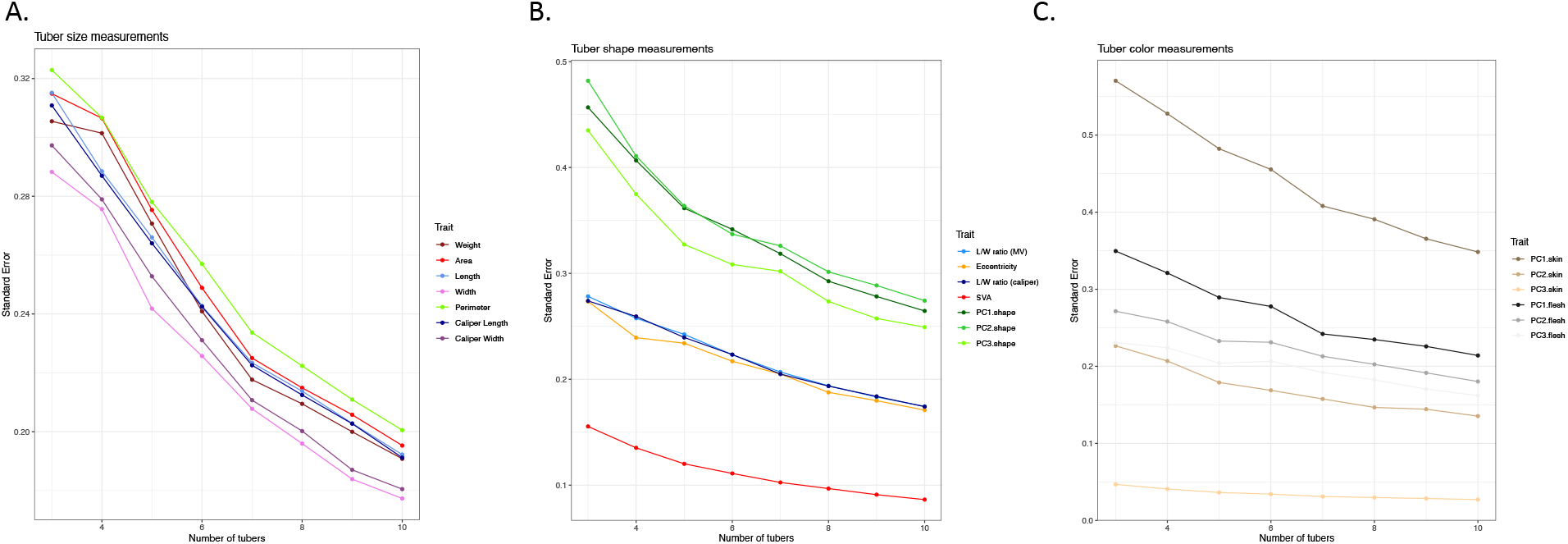
The mean standard errors of trait values given different tuber sample size for each clone within the A08241 breeding population. A) Measurements of tuber size. B) Measurements of tuber shape. C) Measurements of tuber skin and flesh color.

**Fig. S7.**
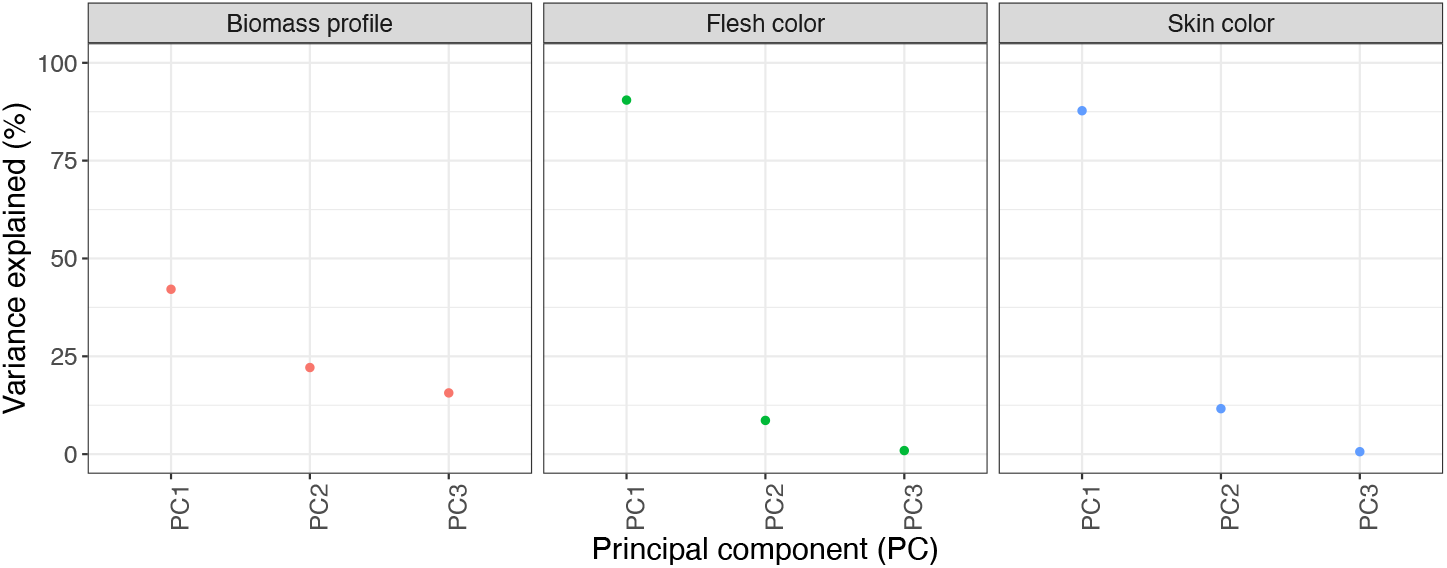
Scree plots illustrating the variance captured by the first 3 principal components for tuber shape, tuber skin, and tuber flesh traits.

**Fig. S8.**
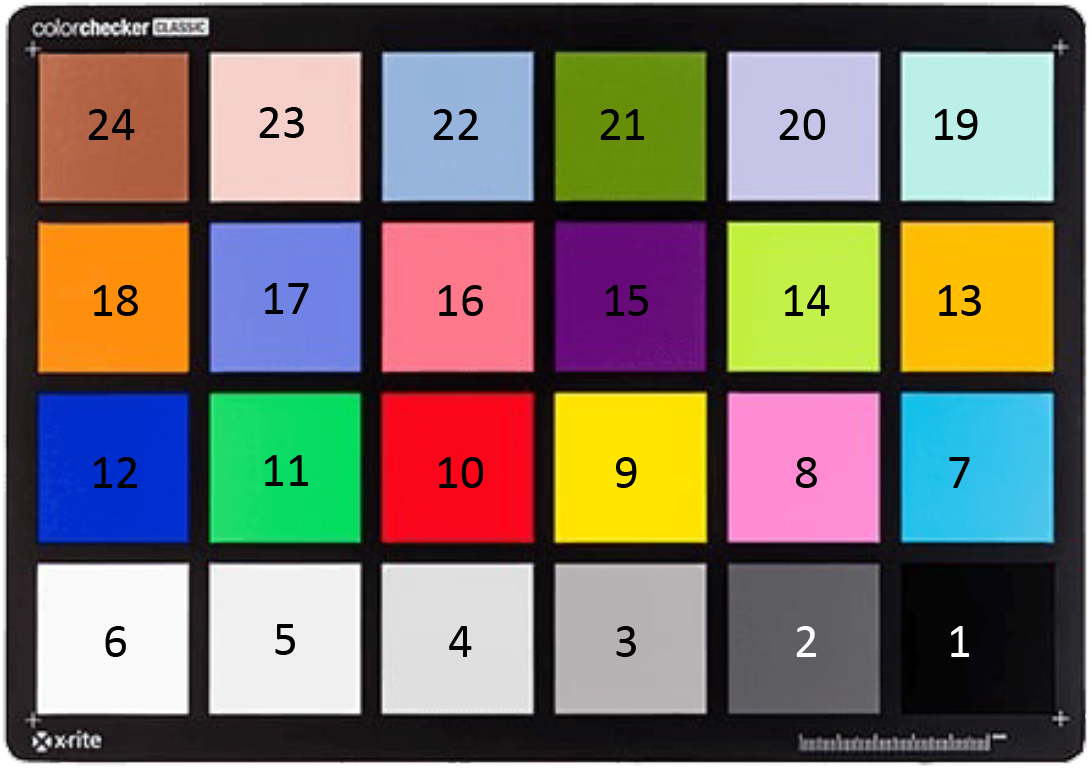
Labeling of color chips on the X-Rite ColorChecker.

**Fig. S9.**
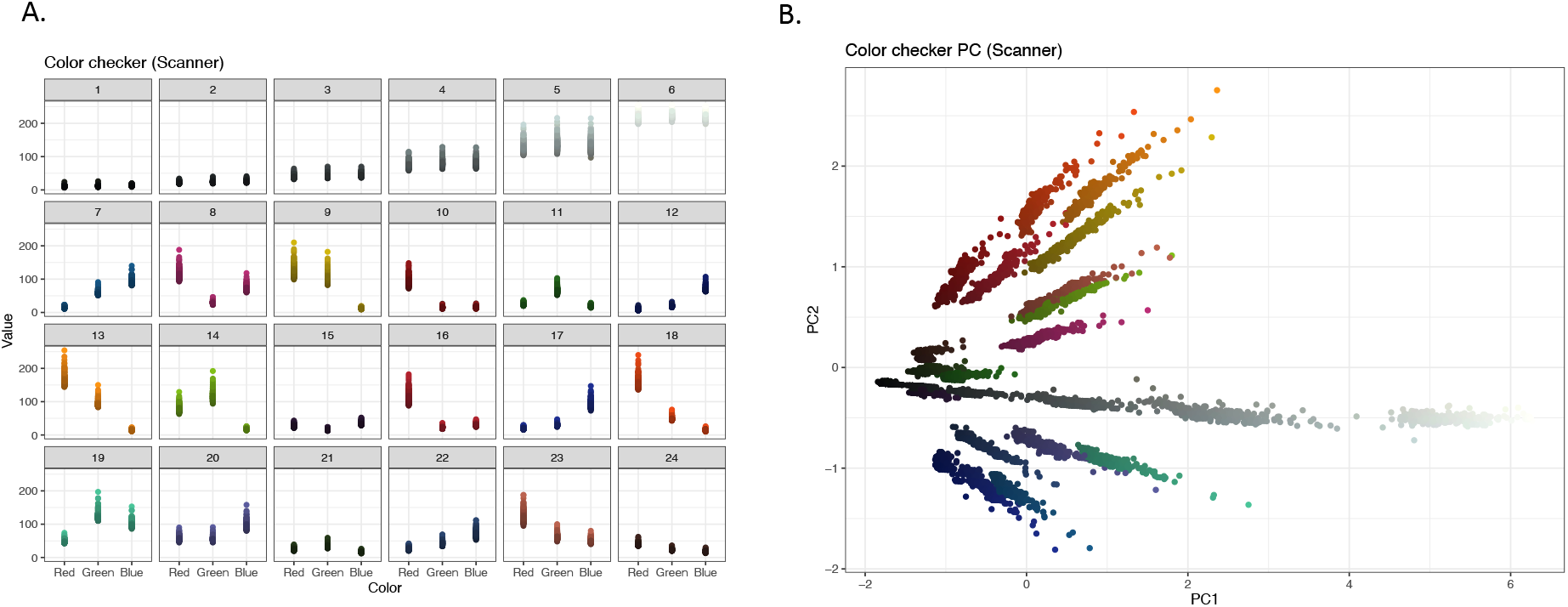
Variation in the color values extracted from chips of a color checker standard on a flatbed scanner. A) Color checker values derived from R, G, and B channels for each of the 24 color checker chips. B) Visualization of color checker values acquired using a flatbed scanner after dimensional reduction principal component analysis.

**Fig. S10.**
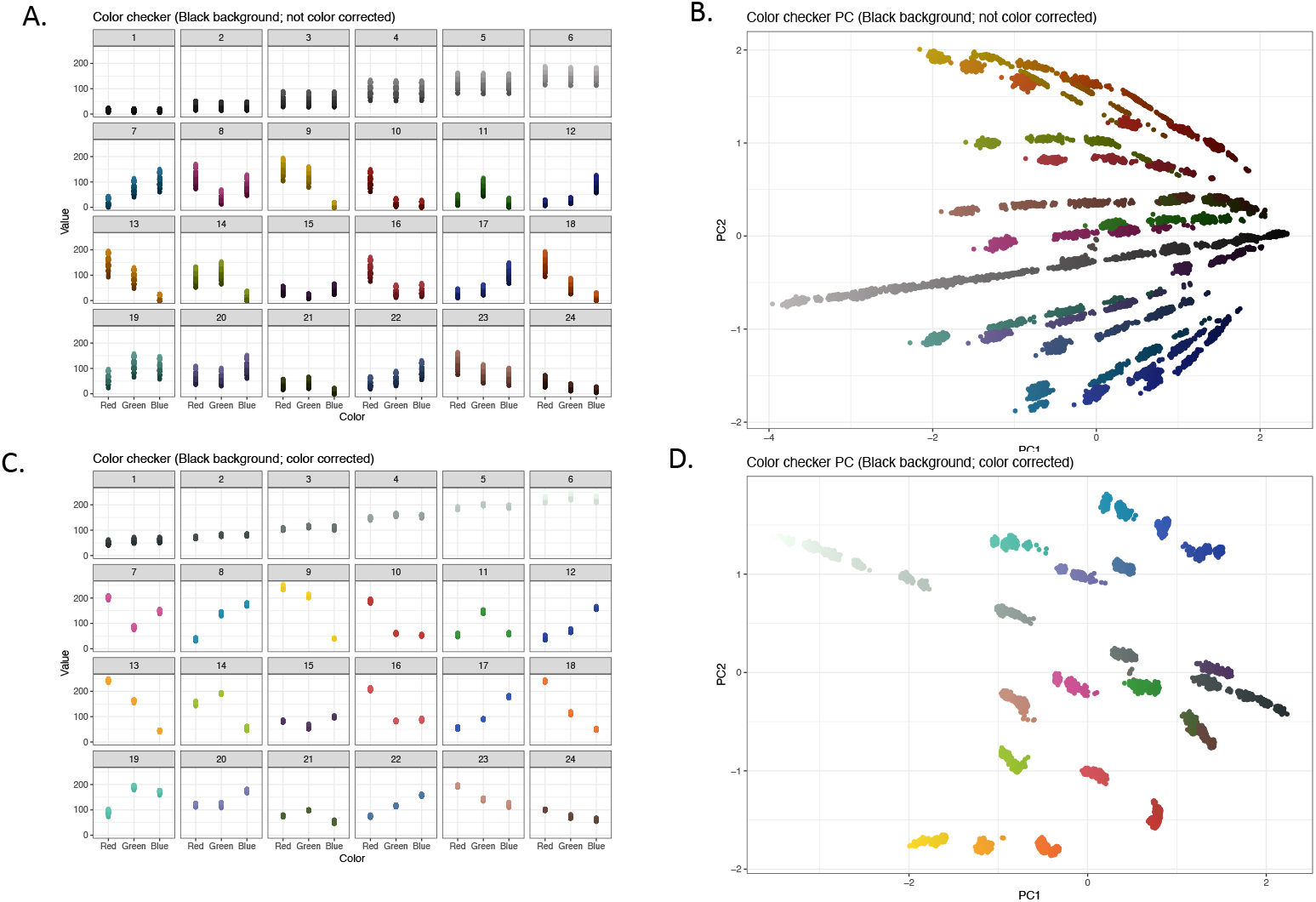
Variation in the color values extracted from chips of a color checker standard using top-down imaging with a RGB camera on a non-reflective black background. A) Color checker values derived from R, G, and B channels for each of the 24 color checker chips without color correction. B) Visualization of non-corrected color checker values after dimensional reduction principal component analysis. C) Color checker values derived from R, G, and B channels for each of the 24 color checker chips after color correction. D) Visualization of color corrected color checker values after dimensional reduction principal component analysis.

**Fig. S11.**
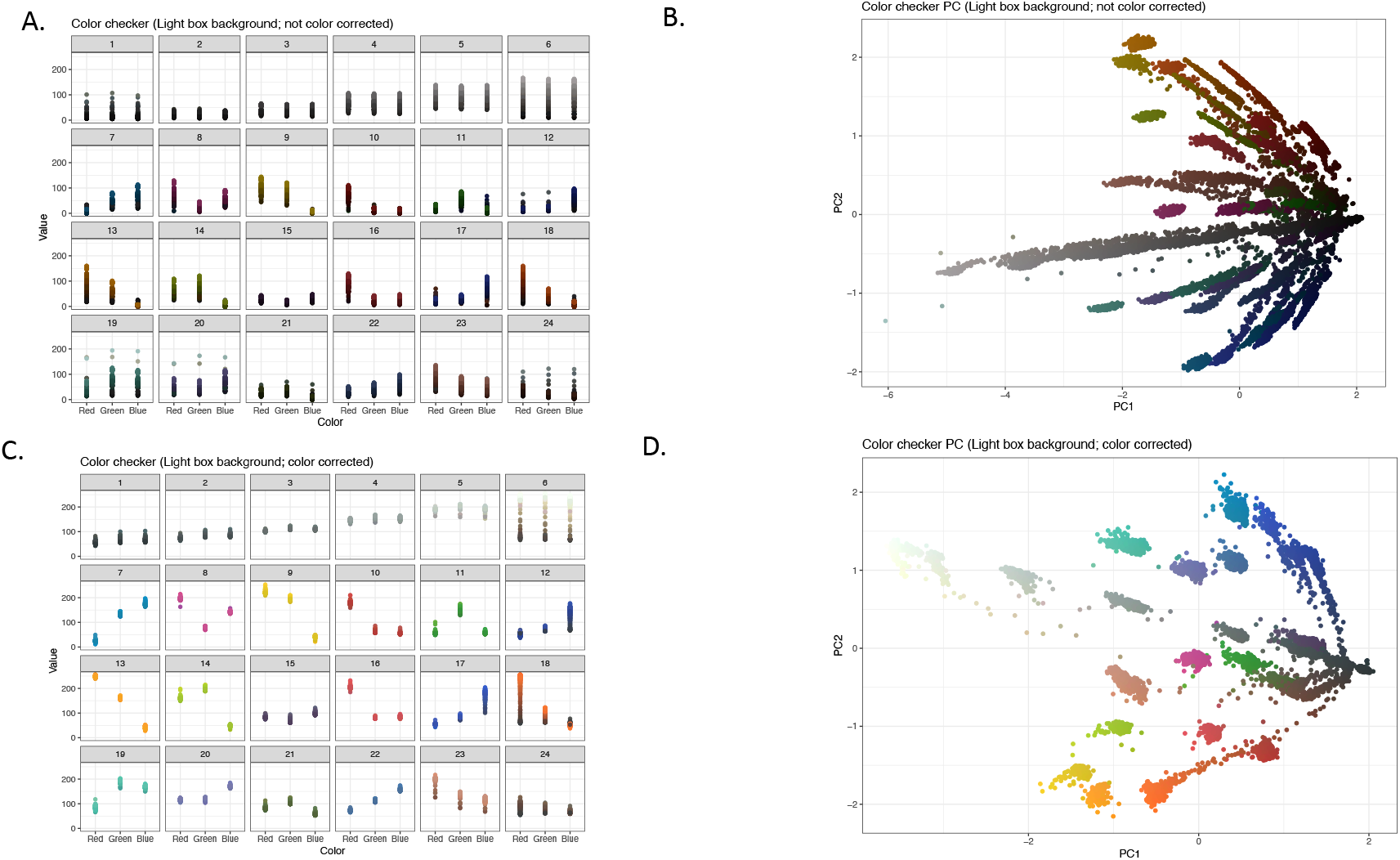
Variation in the color values extracted from chips of a color checker standard using top-down imaging with a RGB camera on a light box background. A) Color checker values derived from R, G, and B channels for each of the 24 color checker chips without color correction. B) Visualization of non-corrected color checker values after dimensional reduction principal component analysis. C) Color checker values derived from R, G, and B channels for each of the 24 color checker chips after color correction. D) Visualization of color corrected color checker values after dimensional reduction principal component analysis.

**Fig. S12.**
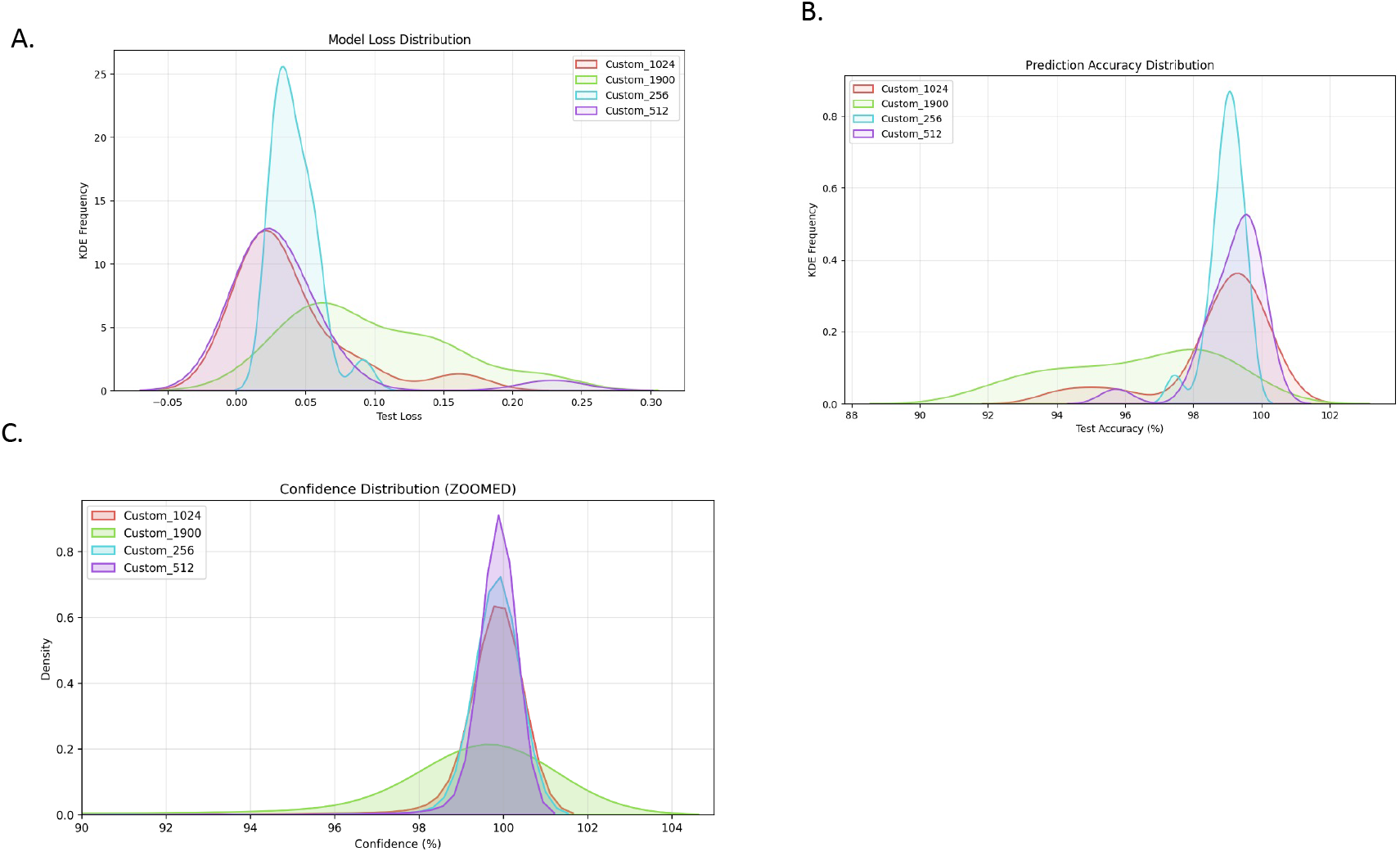
Histograms reporting model performance on the test dataset for images of different resolution. A) Test loss. B) Test accuracy. C) Confidence.

**Fig. S13.**
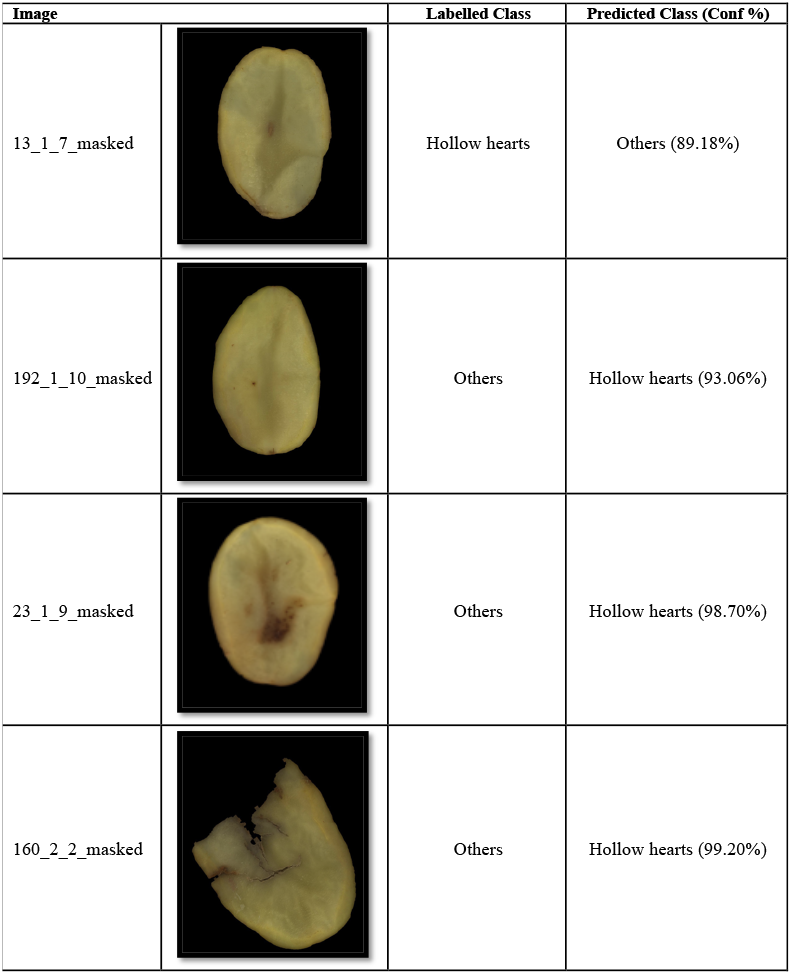
Common misclassifications for less than 100% accurate models.

