## Supplementary material for "A scalable, low-cost phenotyping strategy to assess tuber size, shape, and the colorimetric features of tuber skin and flesh in potato breeding populations": AllFigures

Fig. 1

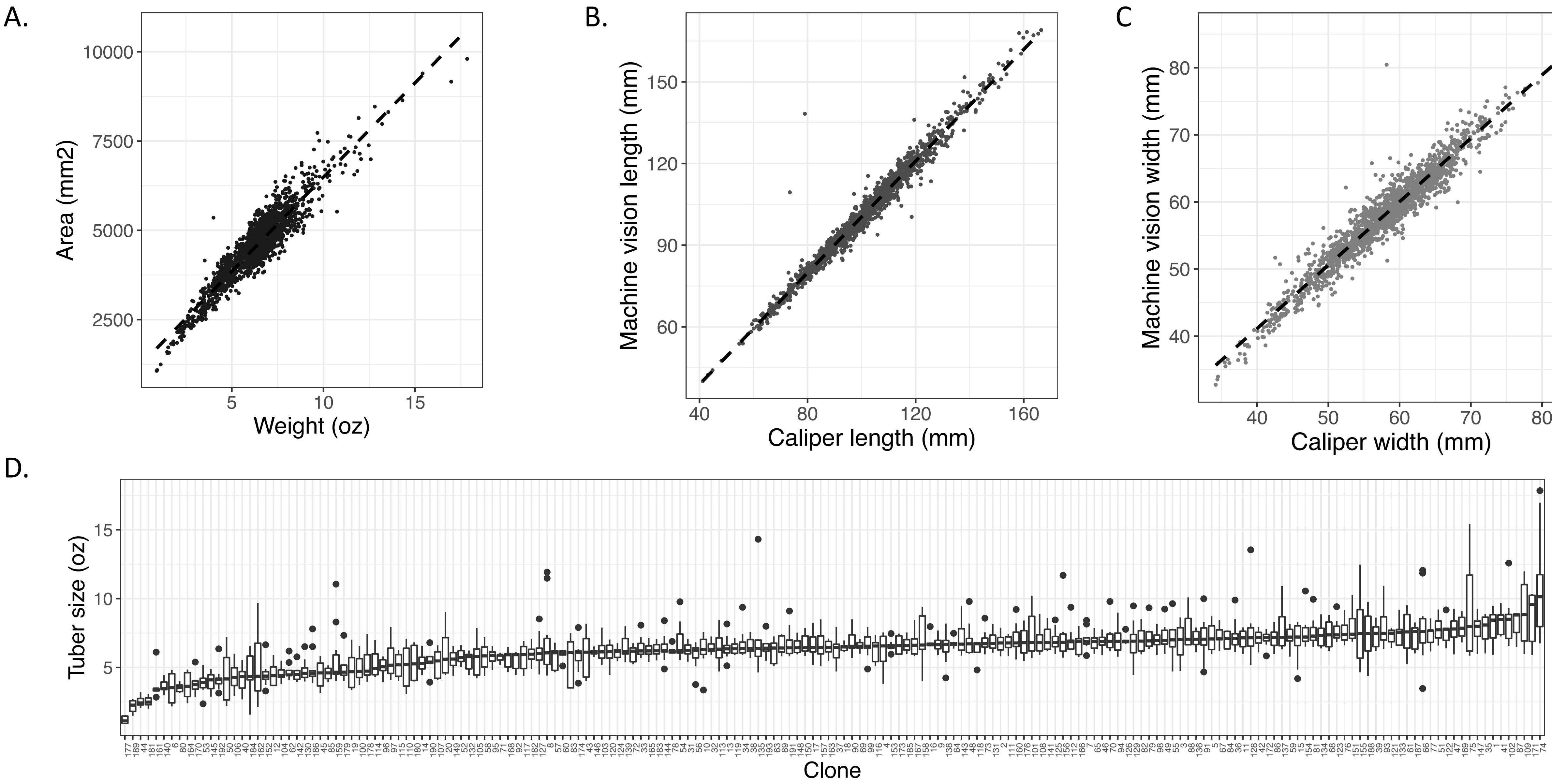

A.
Area (mm2)
10000
7500
5000
2500
5
10
15
Weight (oz)
B.
Machine vision length (mm)
150
120
90
60
40
80
120
160
Caliper length (mm)
C
Machine vision width (mm)
80
70
60
50
40
40
50
60
70
80
Caliper width (mm)
D.
Tuber size (oz)
15
10
5
Clone

Fig. 2

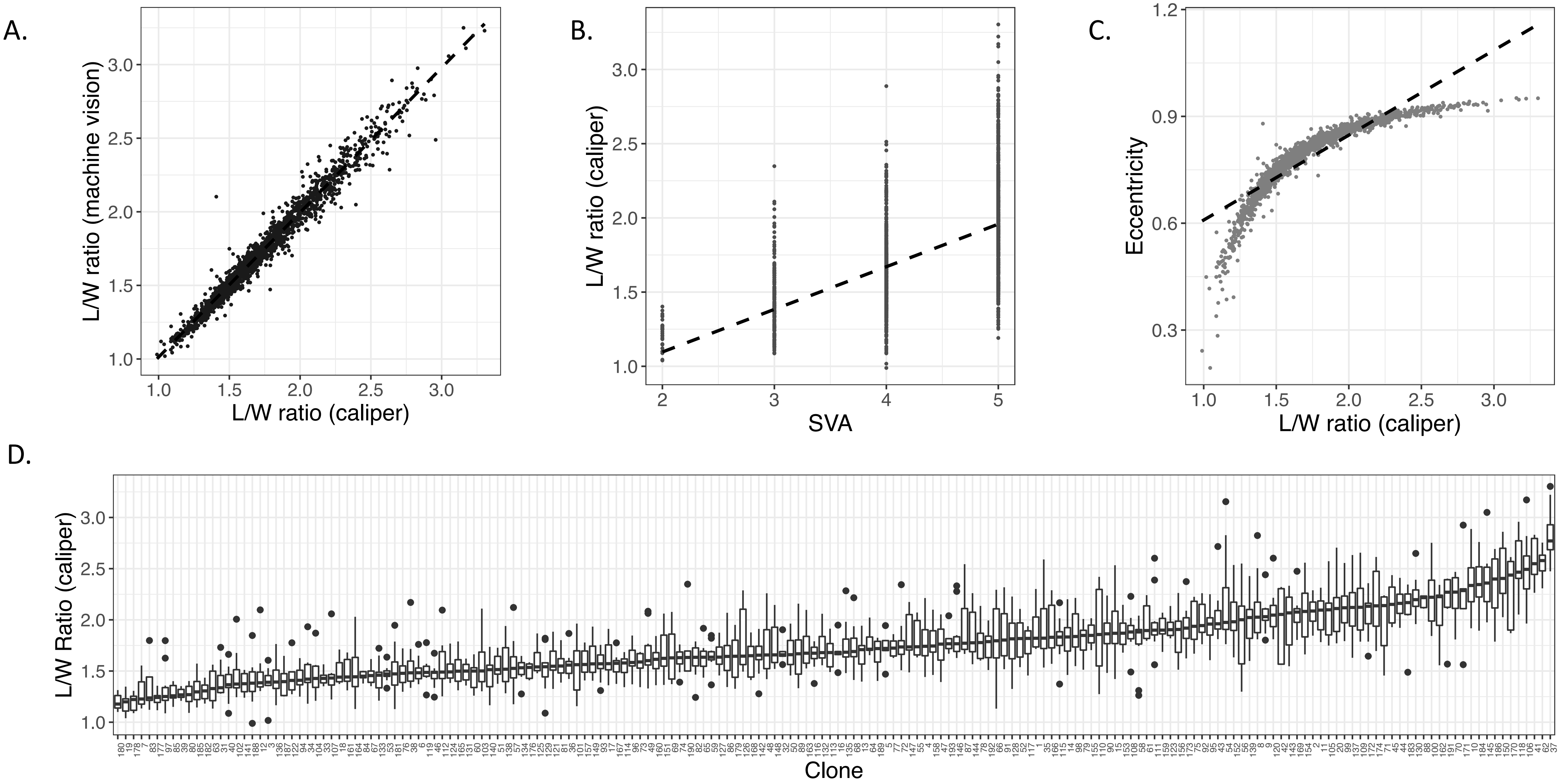
A.
L/W ratio (machine vision)
L/W ratio (caliper)
B.
L/W ratio (caliper)
SVA
C.
Eccentricity
L/W ratio (caliper)
D.
L/W Ratio (caliper)
Clone

Fig. 3

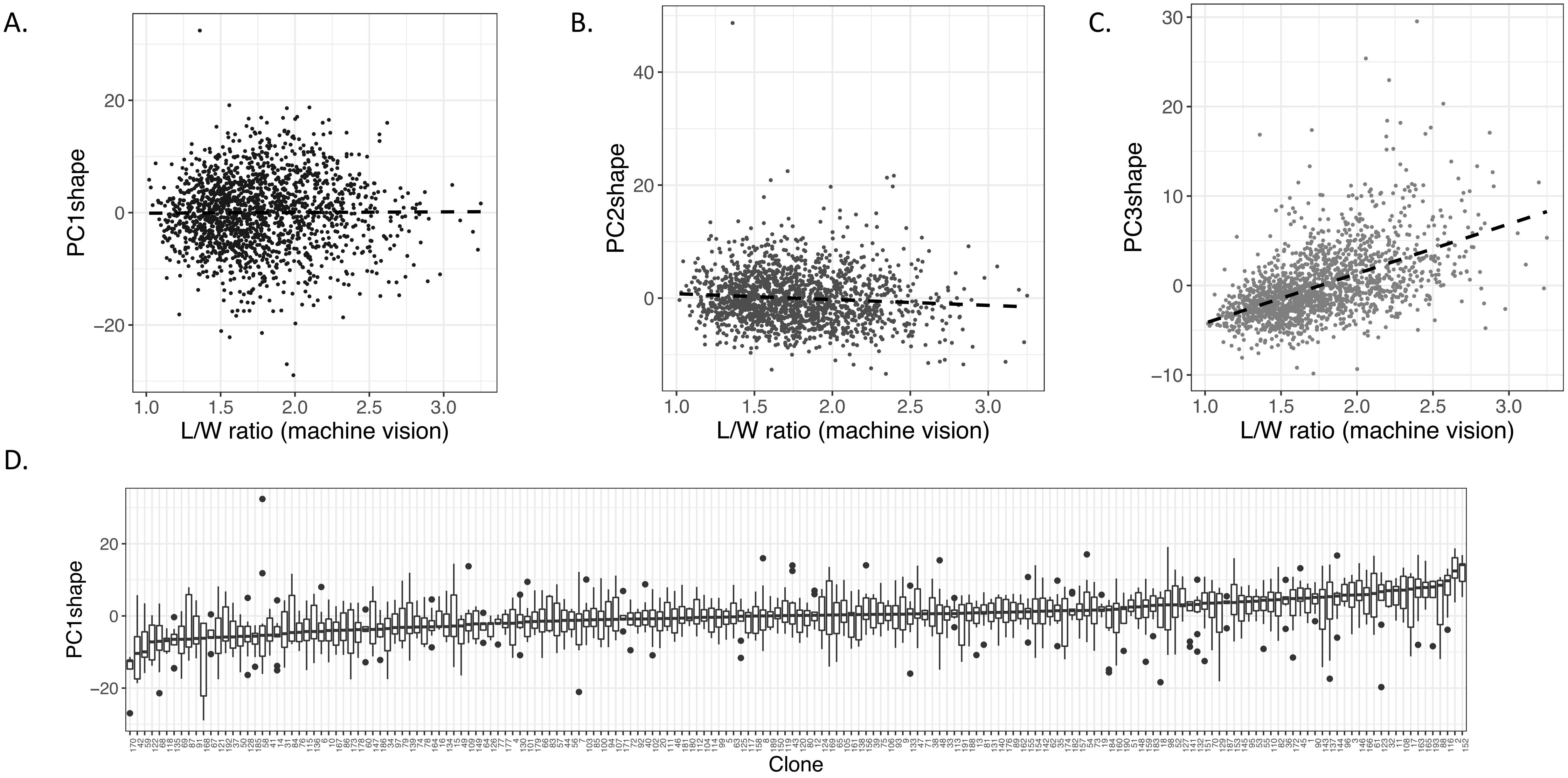
A.
B.
C.
D.
PC1shape
PC2shape
PC3shape
L/W ratio (machine vision)
L/W ratio (machine vision)
L/W ratio (machine vision)
PC1shape
Clone

Fig. 4

A.

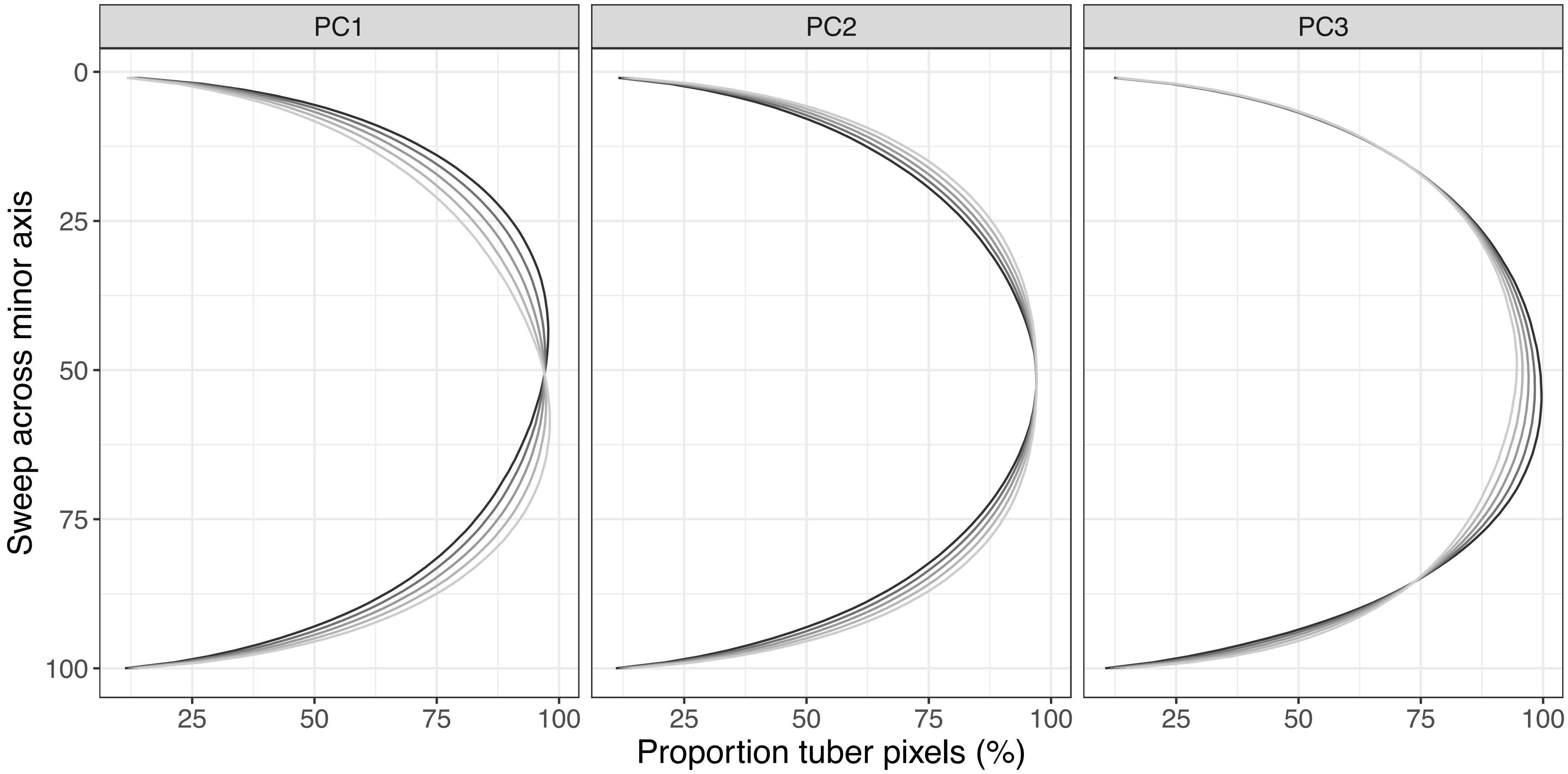

PC1
PC2
PC3
Sweep across minor axis
0
25
50
75
100
25
50
75
100
Proportion tuber pixels (%)

B. Fig. 4 (cont.)

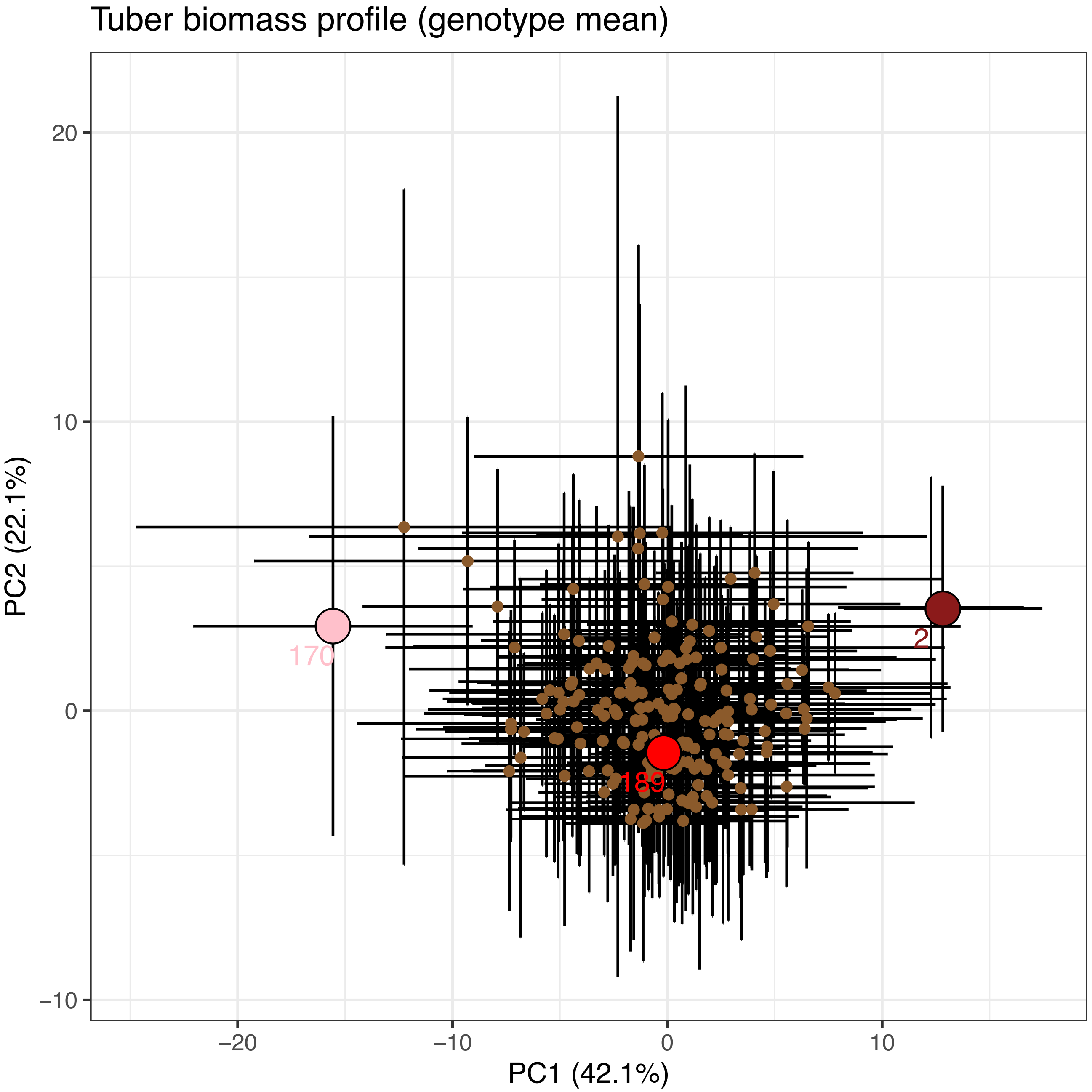

Clone 002

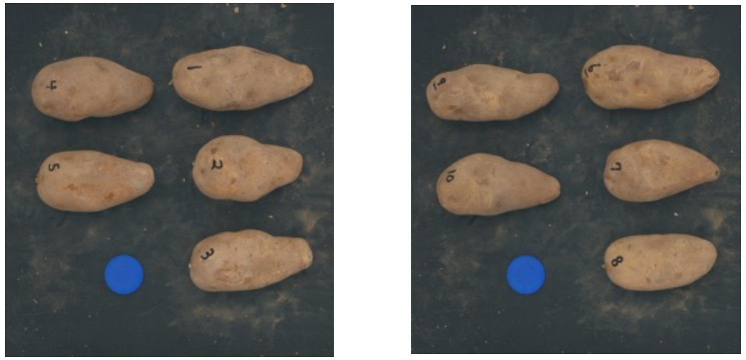

Clone 189

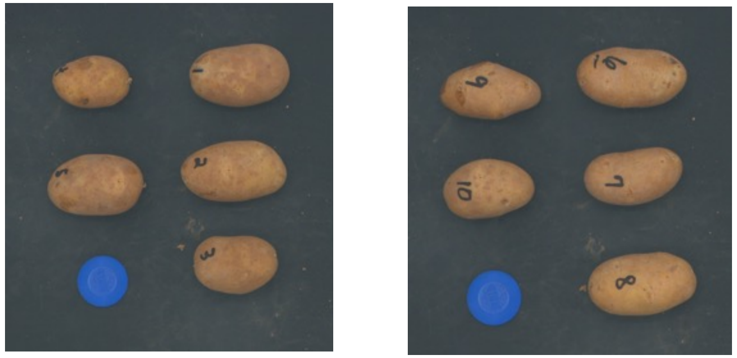

Clone 170

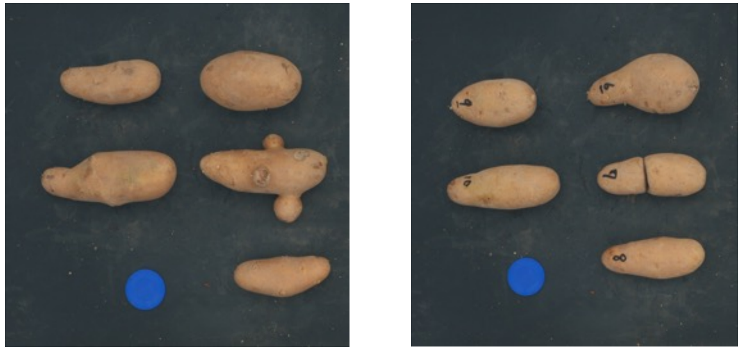

Fig. 5

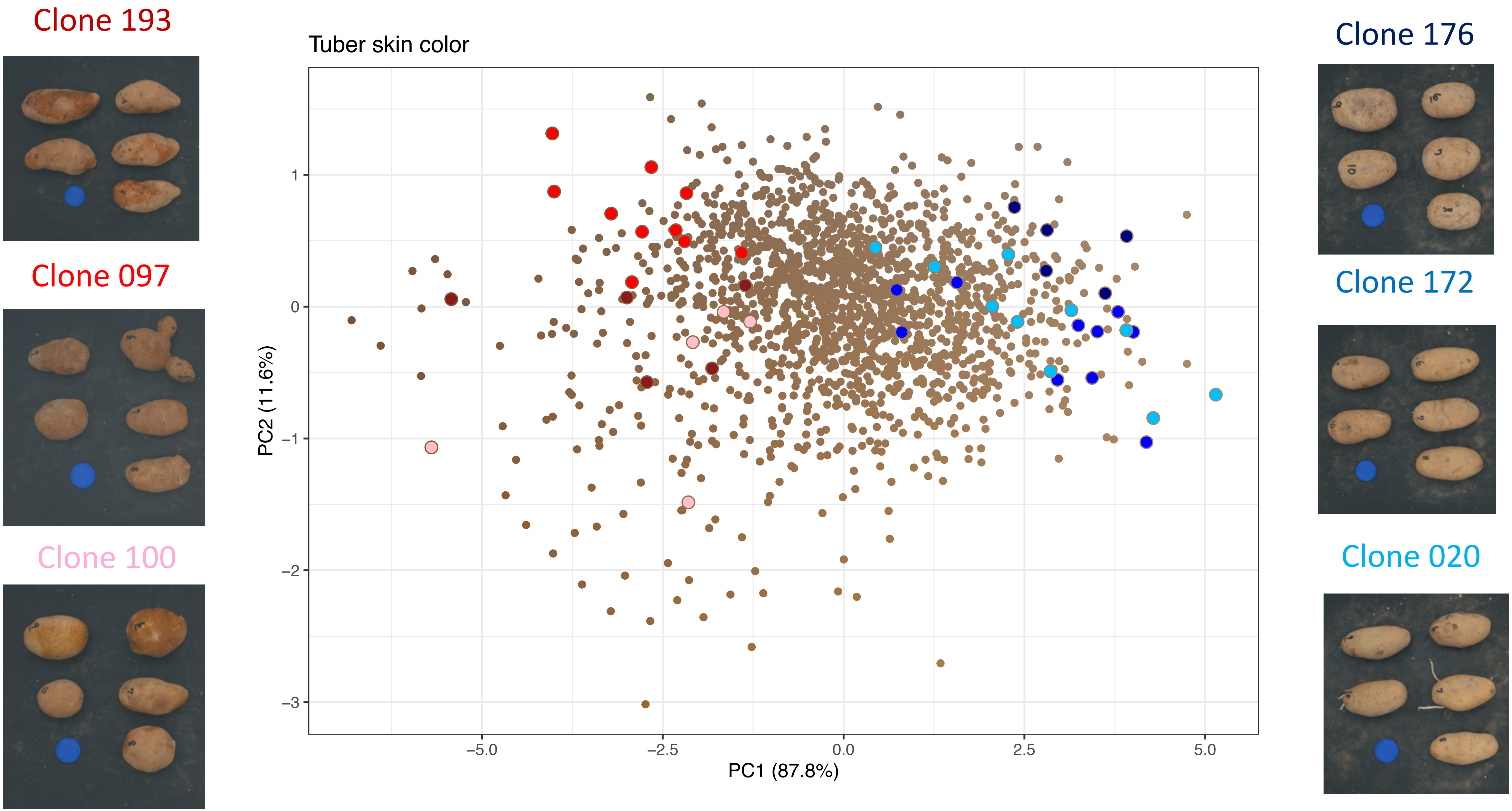
Clone 193
Clone 097
Clone 100
Tuber skin color
1
0
−1
−2
−3
PC2 (11.6%)
−5.0
−2.5
0.0
2.5
5.0
PC1 (87.8%)
Clone 176
Clone 172
Clone 020

Fig. 5 (cont.)

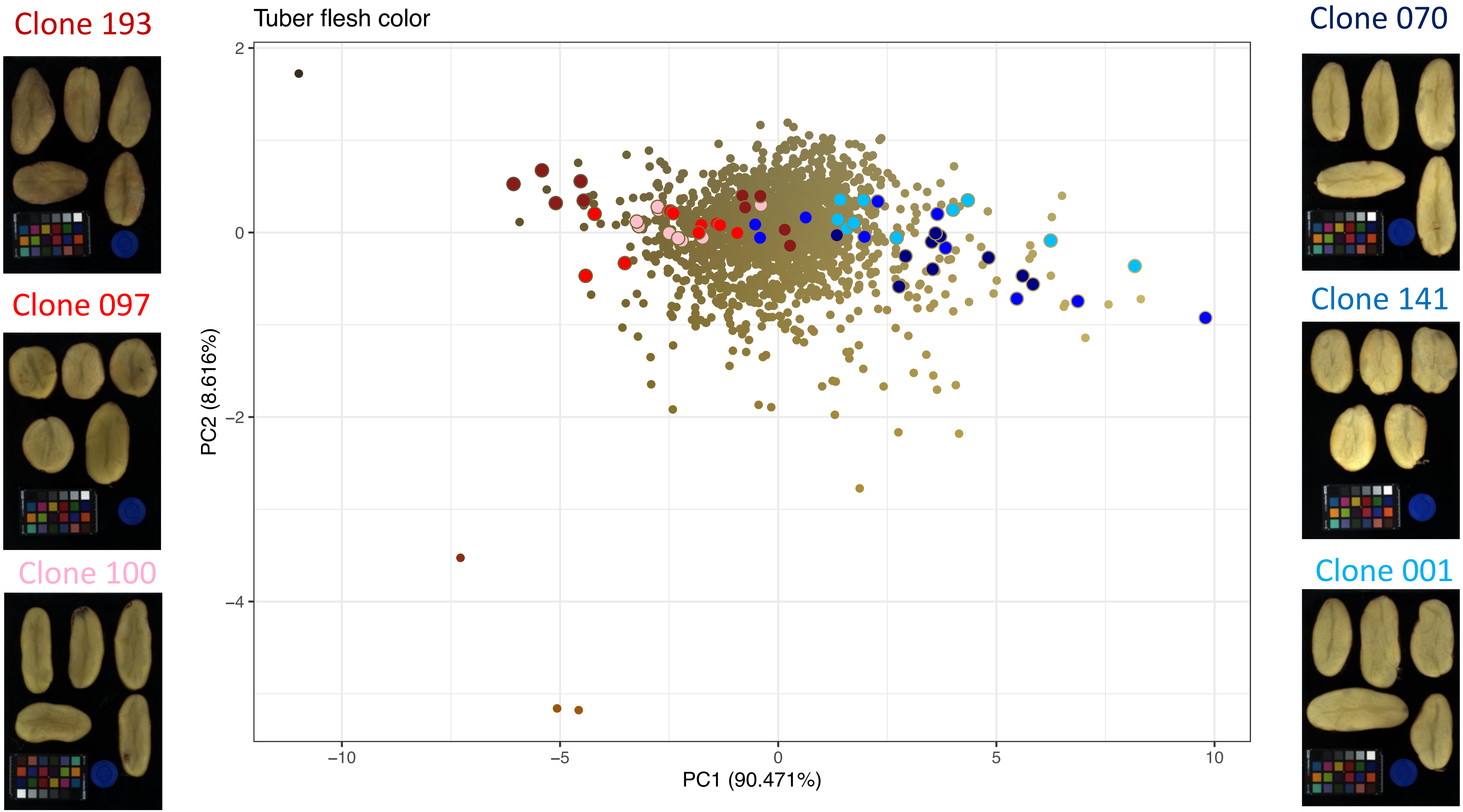
Clone 193
Clone 097
Clone 100
Tuber flesh color
PC2 (8.616%)
PC1 (90.471%)
2
0
−2
−4
−10
−5
0
5
10
Clone 070
Clone 141
Clone 001

Fig. 6

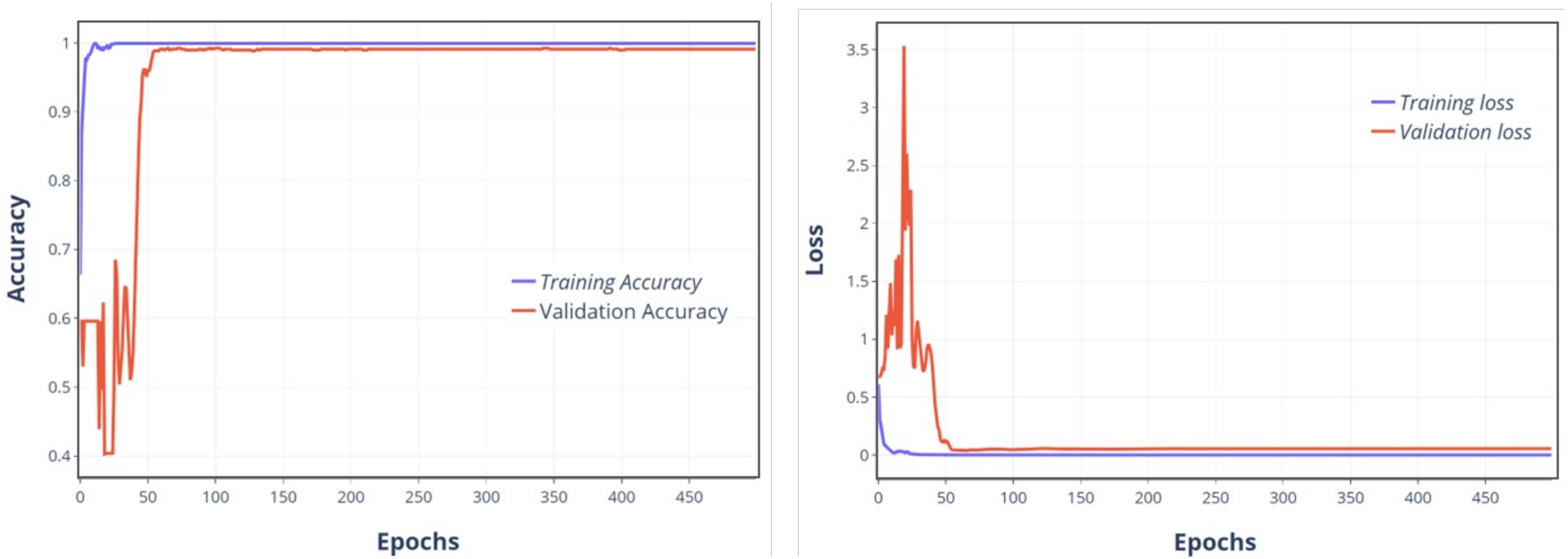

Accuracy
1
0.9
0.8
0.7
0.6
0.5
0.4
0
50
100
150
200
250
300
350
400
450
Epochs
Training Accuracy
Validation Accuracy
Loss
3.5
3
2.5
2
1.5
1
0.5
0
Training loss
Validation loss

Fig. 7

A.

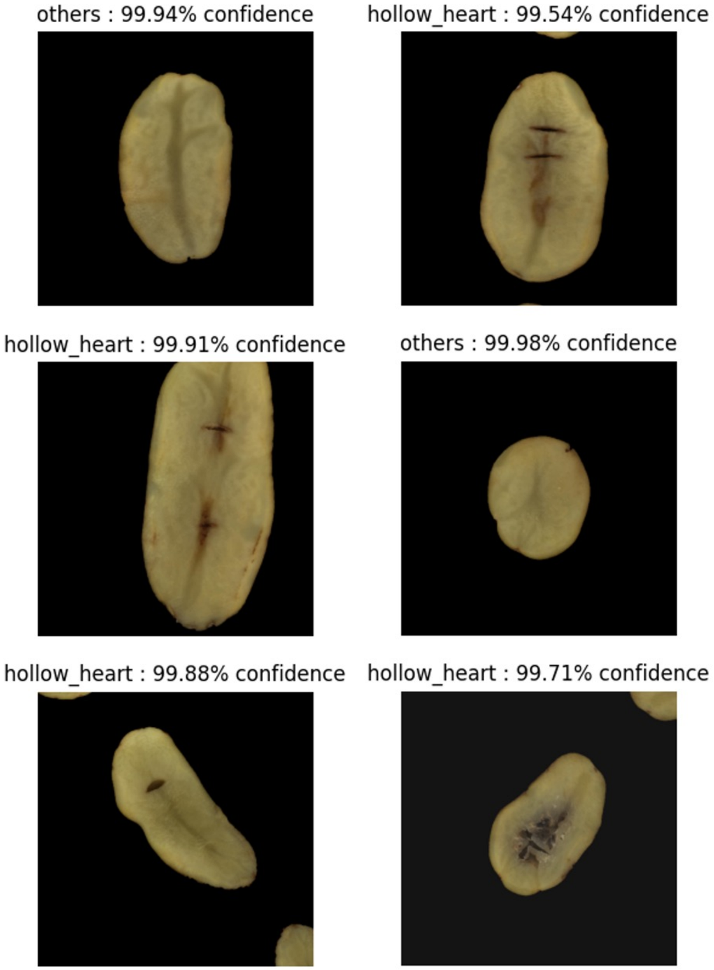

others : 99.94% confidence
hollow_heart : 99.54% confidence
hollow_heart : 99.91% confidence
others : 99.98% confidence
hollow_heart : 99.88% confidence
hollow_heart : 99.71% confidence

B.

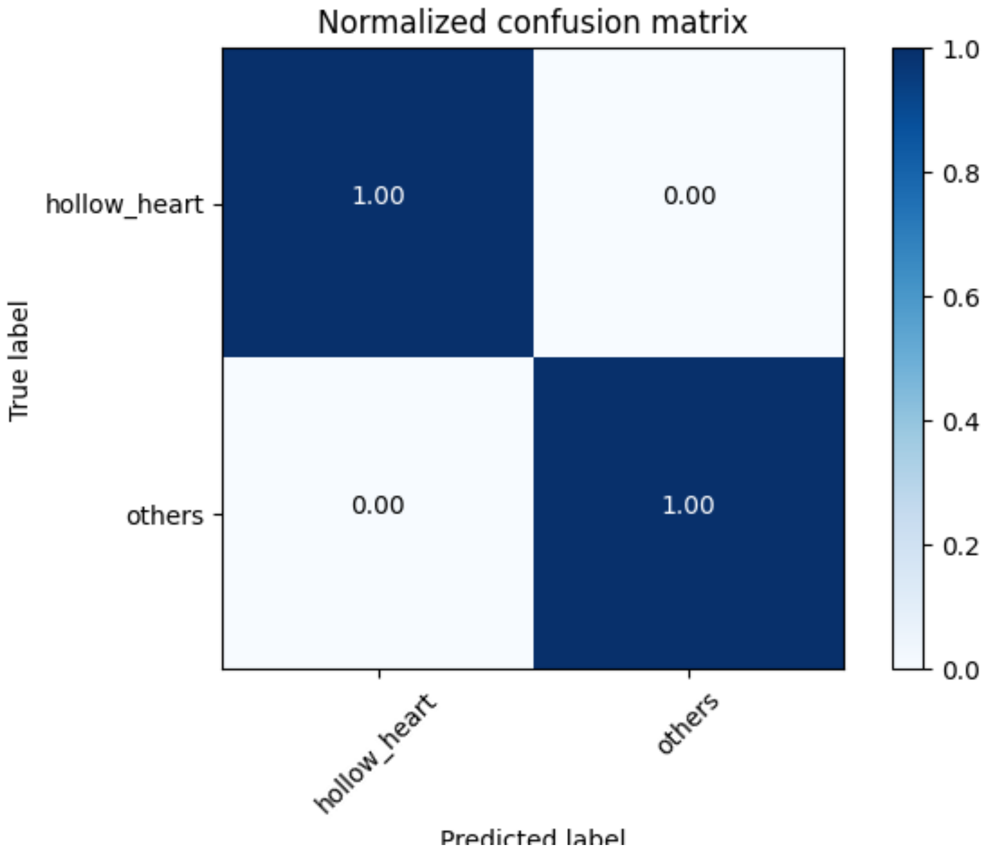

Normalized confusion matrix
1.00
0.00
0.00
1.00
hollow_heart
others
True label
hollow_heart
others
Predicted label
1.0
0.8
0.6
0.4
0.2
0.0

Fig. S1

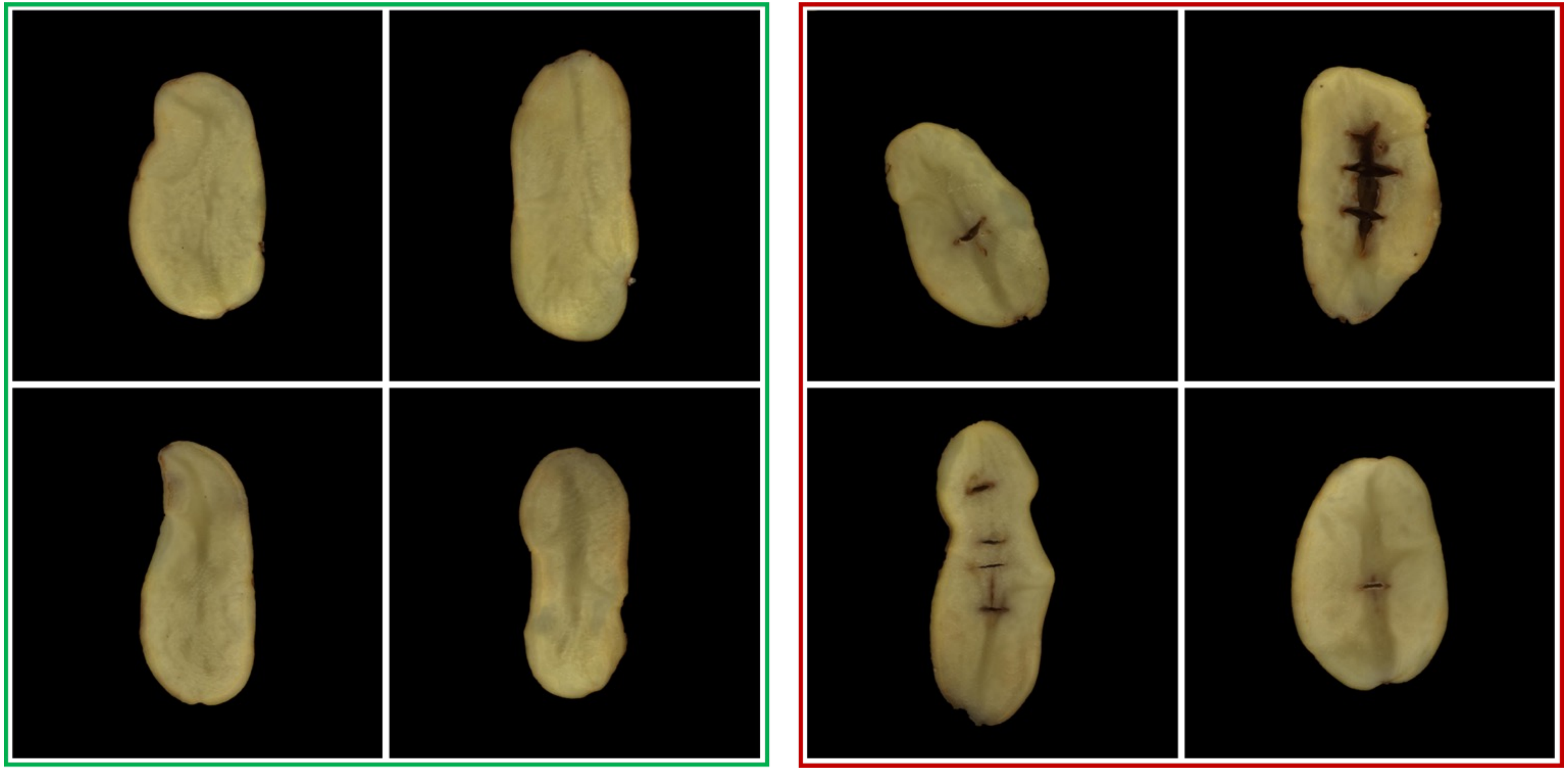

Fig. S2

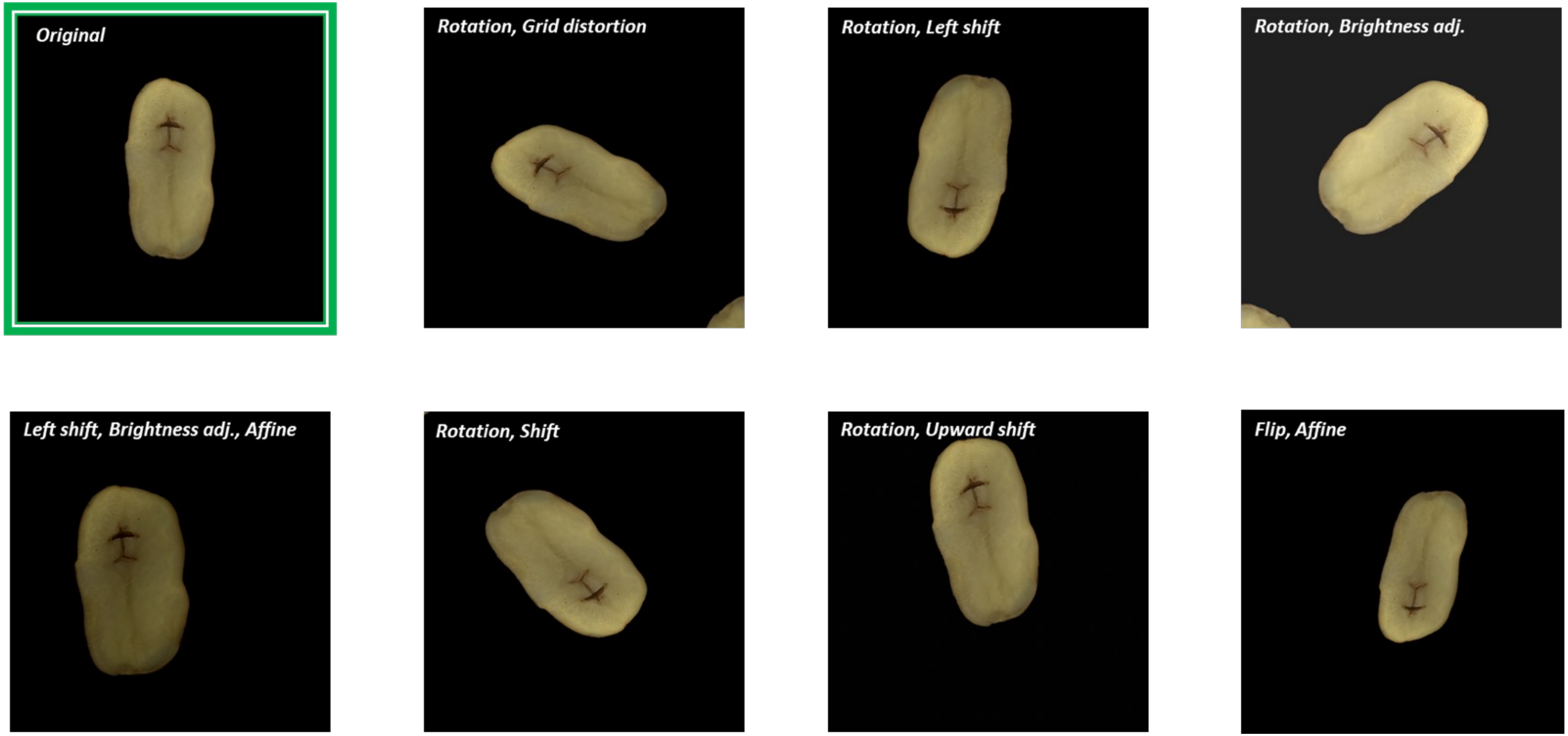
Original
Rotation, Grid distortion
Rotation, Left shift
Rotation, Brightness adj.
Left shift, Brightness adj., Affine
Rotation, Shift
Rotation, Upward shift
Flip, Affine

Fig. S3

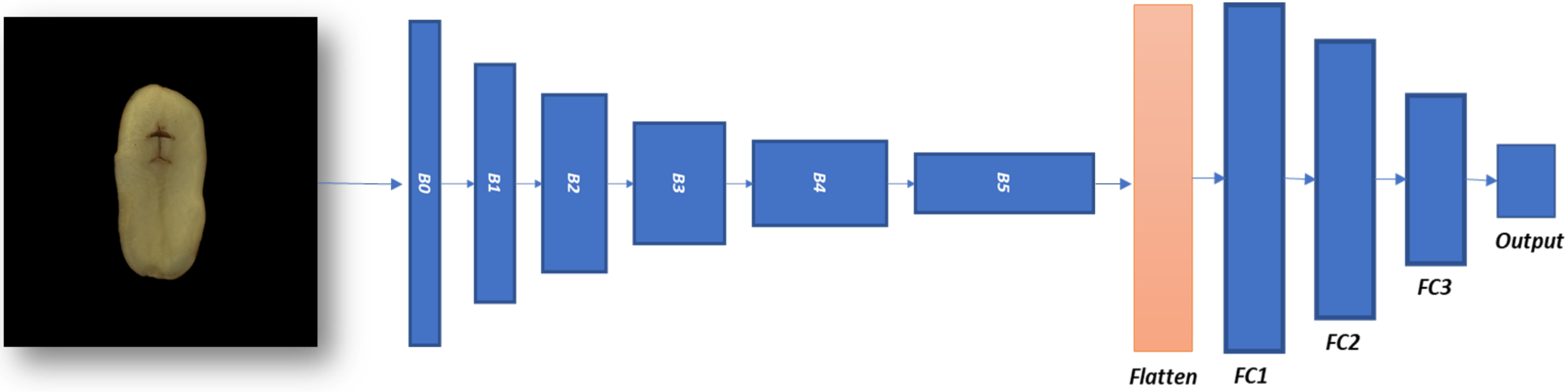

Fig. S4

A. B. C.

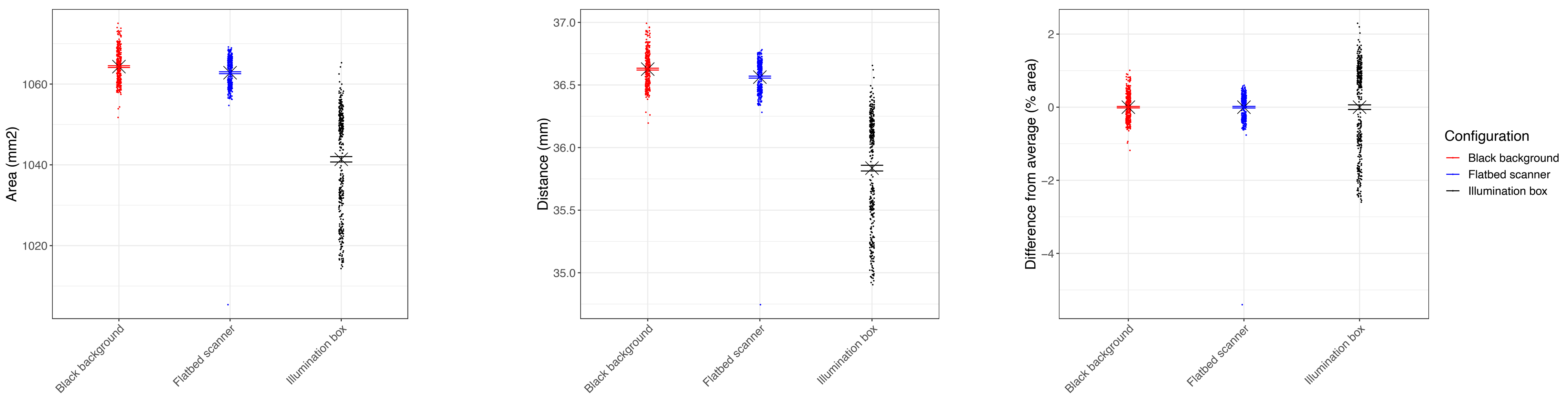
Area (mm2)
1060
1040
1020
Distance (mm)
37.0
36.5
36.0
35.5
35.0
Difference from average (% area)
2
0
−2
−4
Black background
Flatbed scanner
Illumination box
Configuration
Black background
Flatbed scanner
Illumination box

Fig. S5

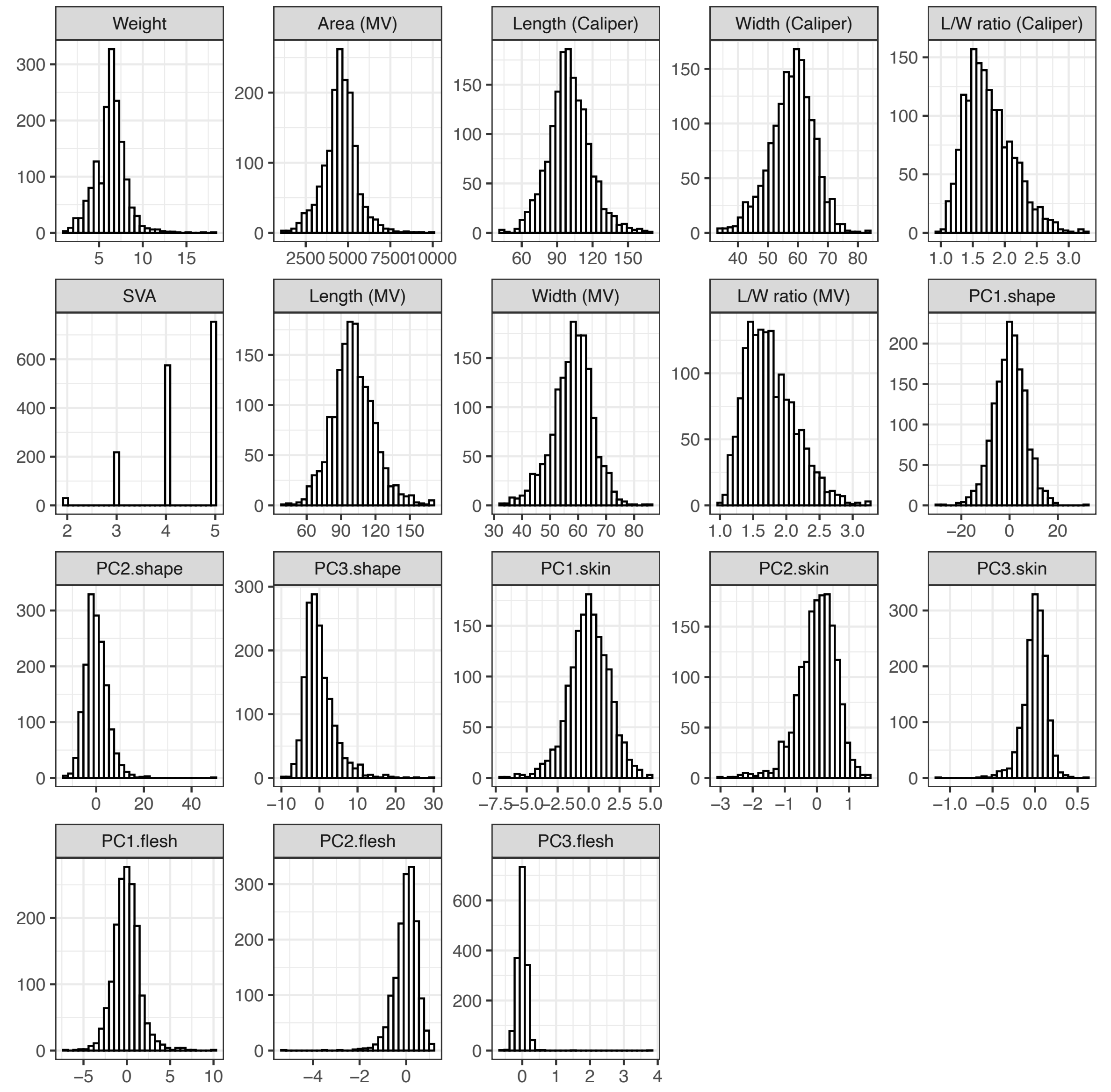
Weight
Area (MV)
Length (Caliper)
Width (Caliper)
L/W ratio (Caliper)
SVA
Length (MV)
Width (MV)
L/W ratio (MV)
PC1.shape
PC2.shape
PC3.shape
PC1.skin
PC2.skin
PC3.skin
PC1.flesh
PC2.flesh
PC3.flesh

Fig. S6

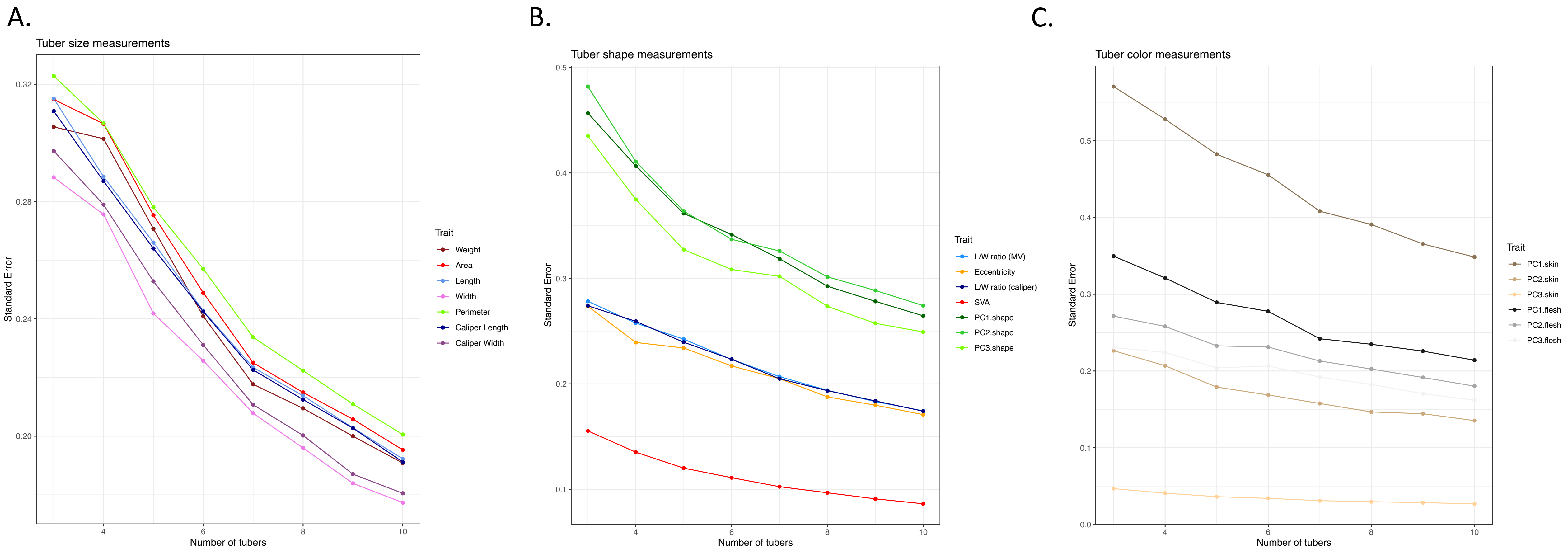

A.
Tuber size measurements
Standard Error
Number of tubers
Trait
Weight
Area
Length
Width
Perimeter
Caliper Length
Caliper Width
B.
Tuber shape measurements
Standard Error
Number of tubers
Trait
L/W ratio (MV)
Eccentricity
L/W ratio (caliper)
SVA
PC1.shape
PC2.shape
PC3.shape
C.
Tuber color measurements
Standard Error
Number of tubers
Trait
PC1.skin
PC2.skin
PC3.skin
PC1.flesh
PC2.flesh
PC3.flesh

Fig. S7

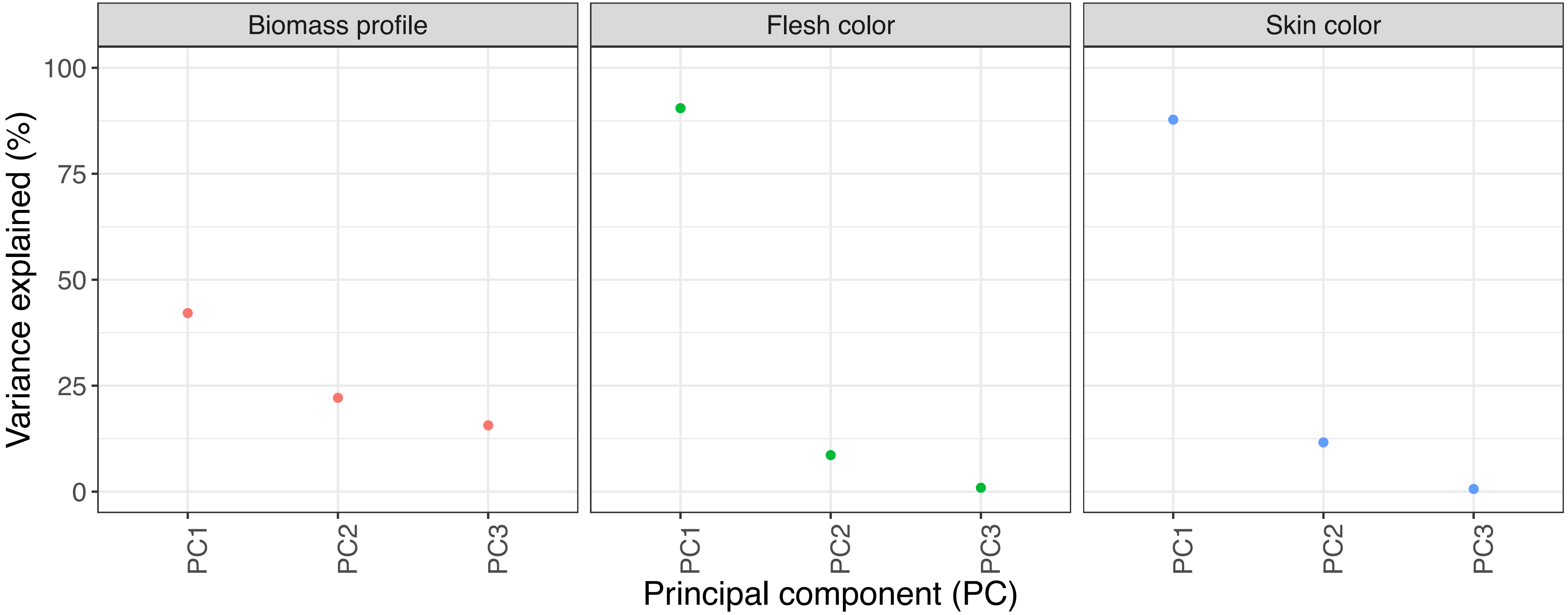

Biomass profile
Flesh color
Skin color
Variance explained (%)
100
75
50
25
0
PC1
PC2
PC3
PC1
PC2
PC3
PC1
PC2
PC3
Principal component (PC)

Fig. S8

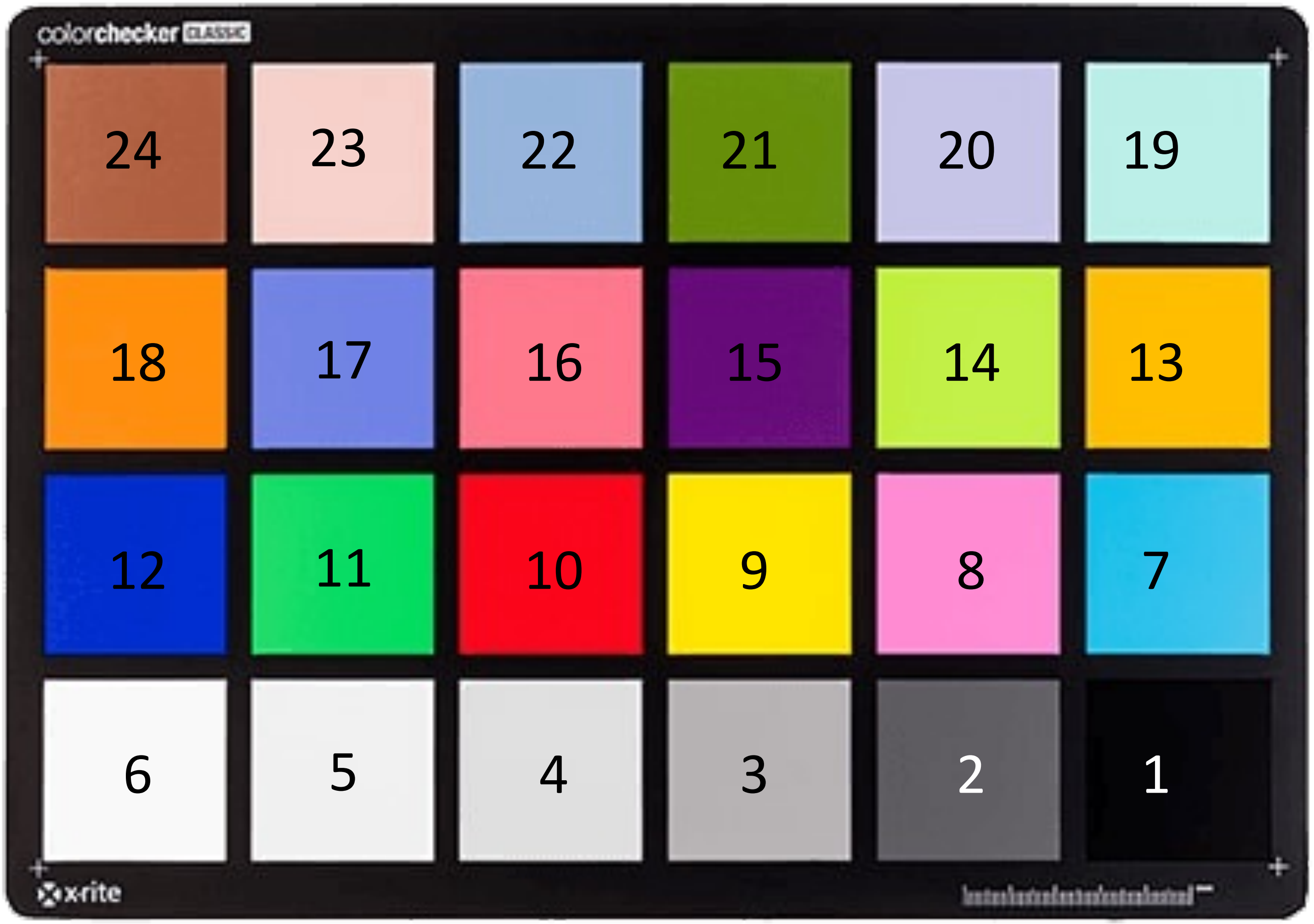
colorchecker CLASSIC
24
23
22
21
20
19
18
17
16
15
14
13
12
11
10
9
8
7
6
5
4
3
2
1
x-rite

Fig. S9

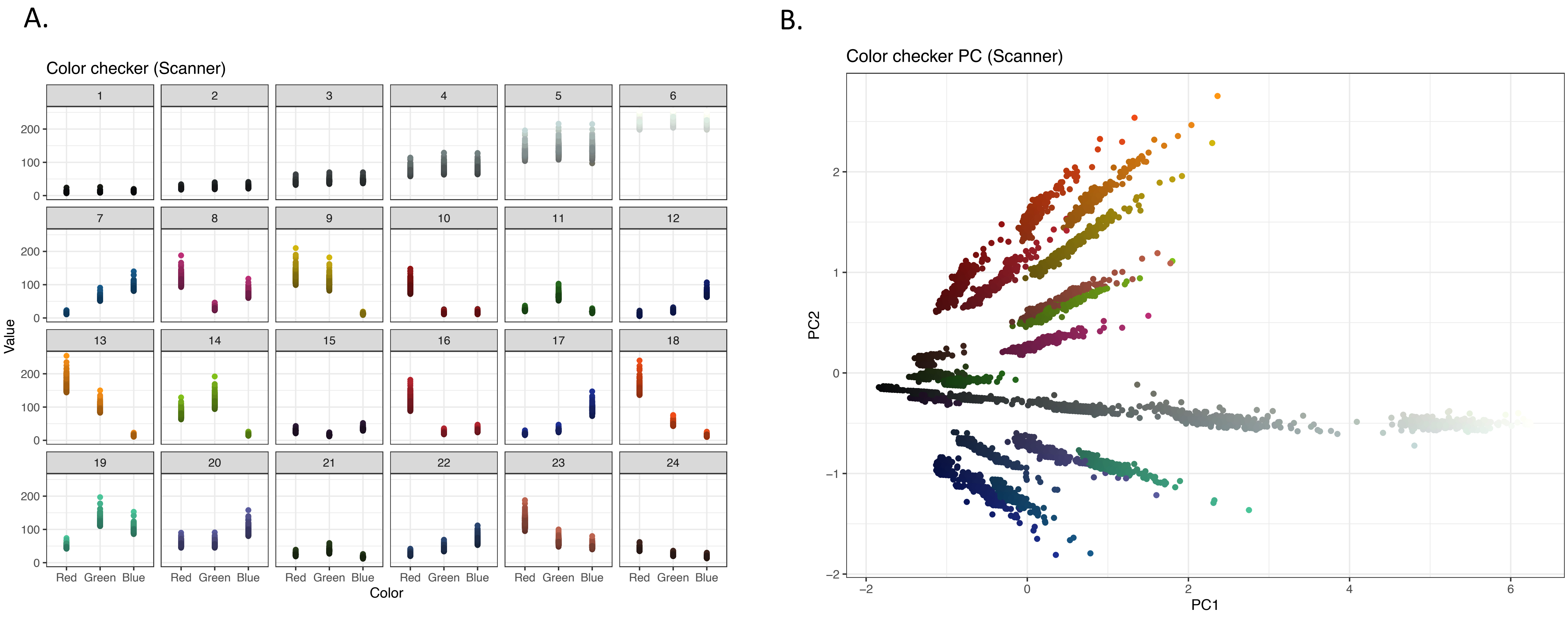
A.
Color checker (Scanner)
Value
0
100
200
Red Green Blue
Color
B.
Color checker PC (Scanner)
PC2
PC1
−2
−1
0
1
2
−2
0
2
4
6

Fig. S10

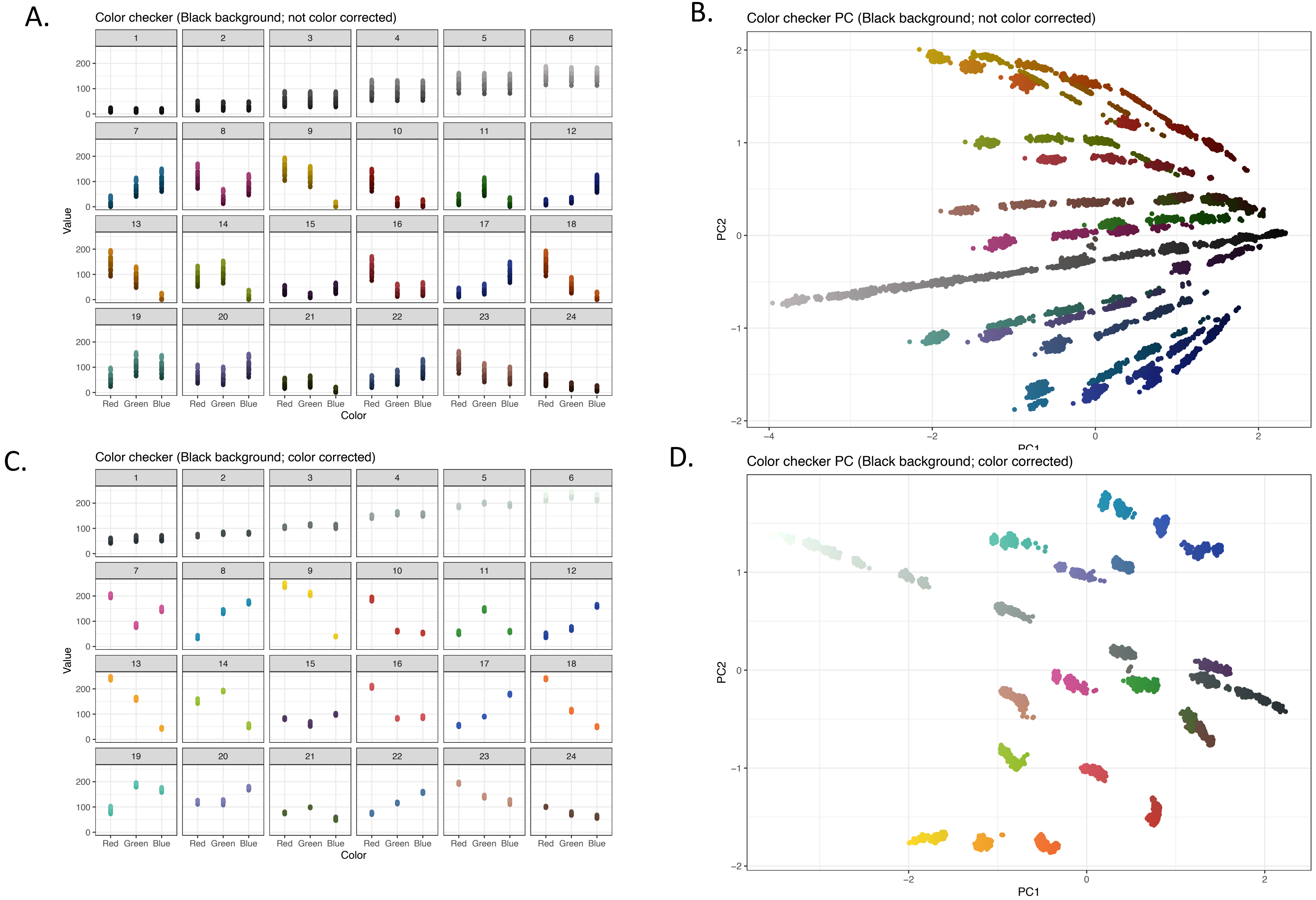
A.
Color checker (Black background; not color corrected)
B.
Color checker PC (Black background; not color corrected)
C.
Color checker (Black background; color corrected)
D.
Color checker PC (Black background; color corrected)
Value
Color
Red Green Blue
PC1
PC2

Fig. S11

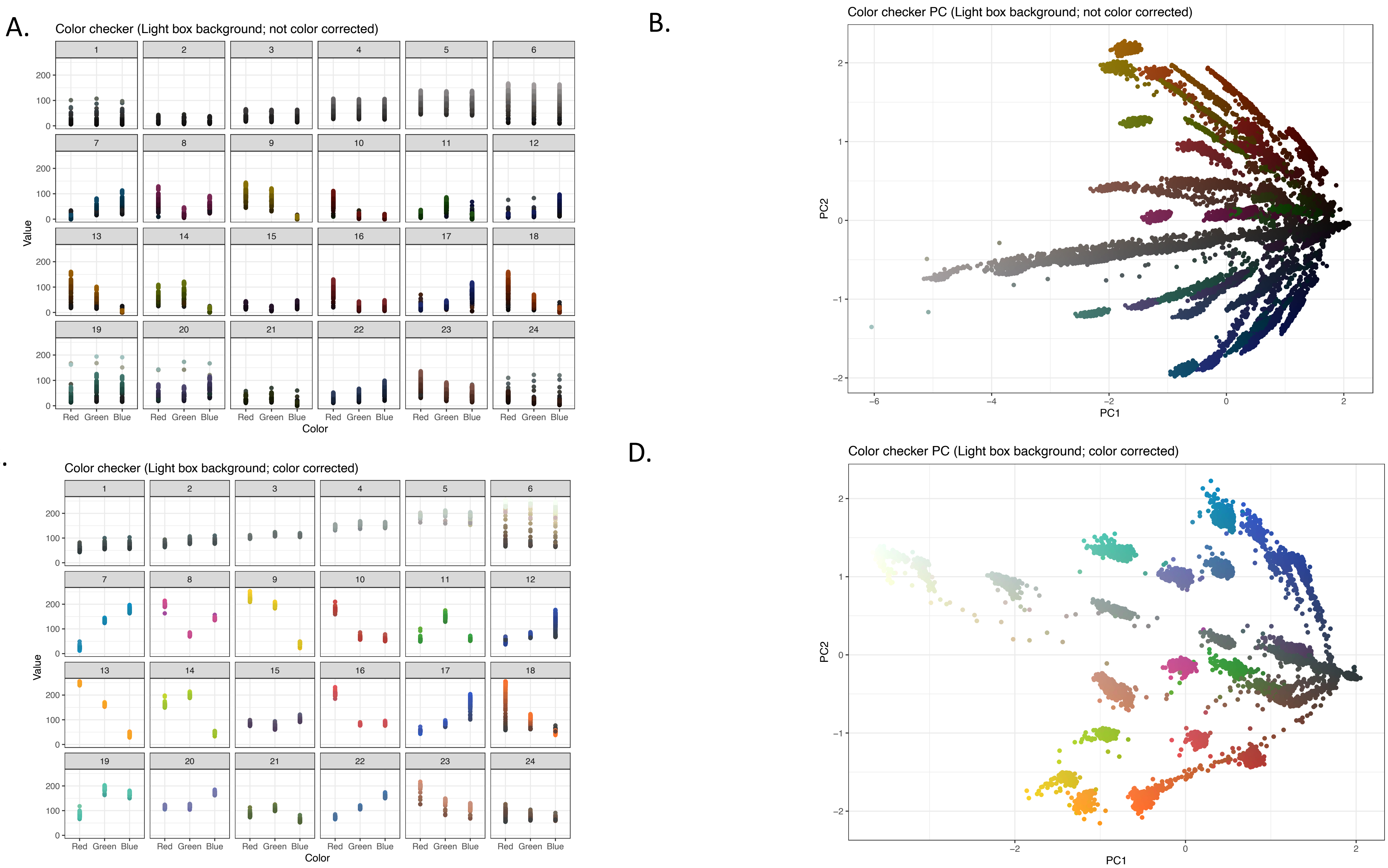
A.
Color checker (Light box background; not color corrected)
Value
Color
Red Green Blue
B.
Color checker PC (Light box background; not color corrected)
PC1
PC2
C.
Color checker (Light box background; color corrected)
Value
Color
Red Green Blue
D.
Color checker PC (Light box background; color corrected)
PC1
PC2

Fig. S12

A.

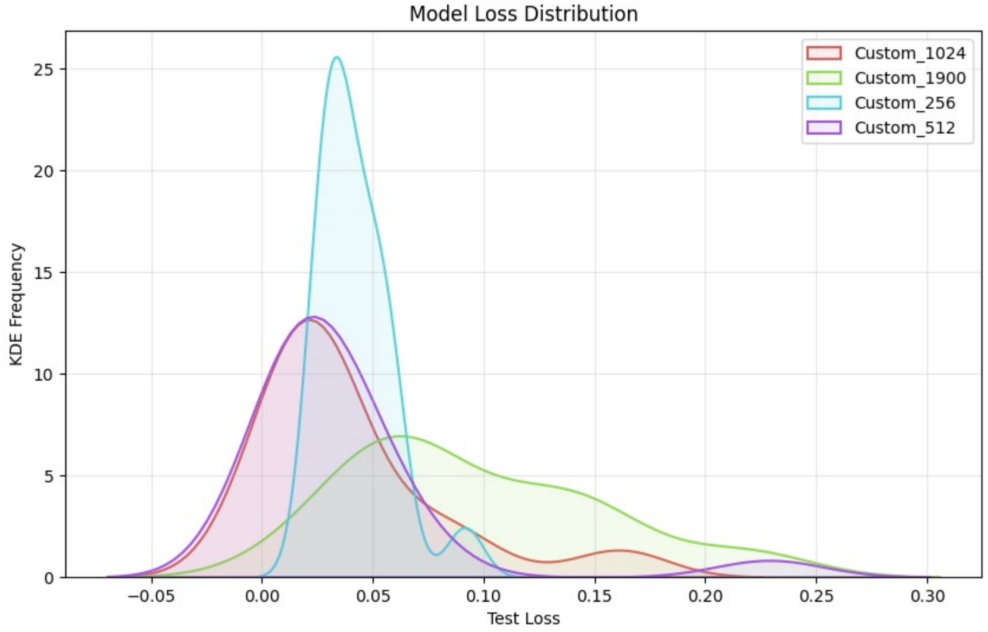
Model Loss Distribution
Custom_1024
Custom_1900
Custom_256
Custom_512
KDE Frequency
Test Loss

B.

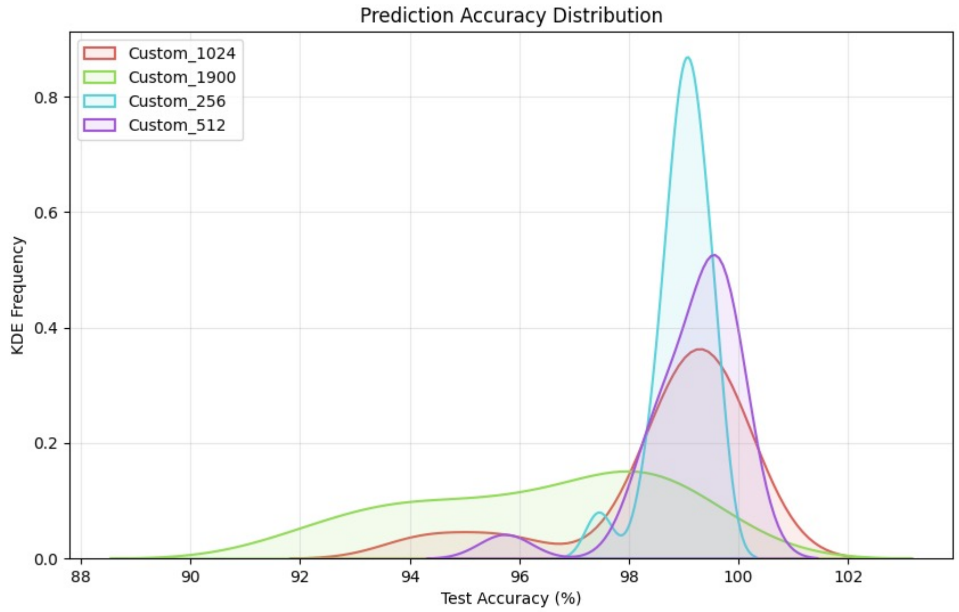
Prediction Accuracy Distribution
Custom_1024
Custom_1900
Custom_256
Custom_512
KDE Frequency
Test Accuracy (%)

C.

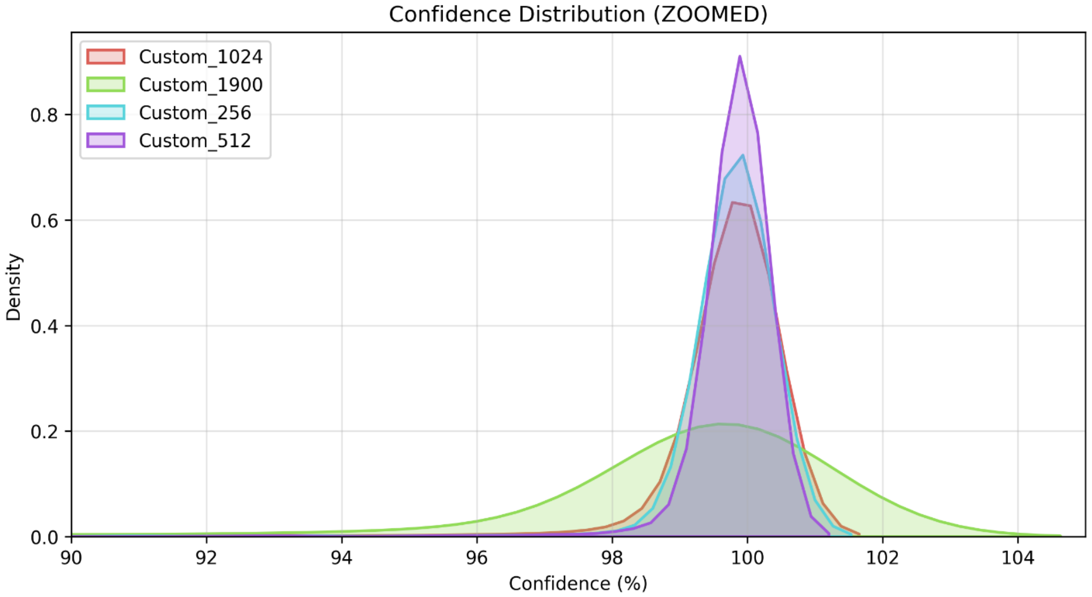
Confidence Distribution (ZOOMED)
Custom_1024
Custom_1900
Custom_256
Custom_512
Density
Confidence (%)

Fig. S13

| **Image** | | **Labelled Class** | **Predicted Class (Conf %)** |
|---|---|---|---|
| 13_1_7_masked | 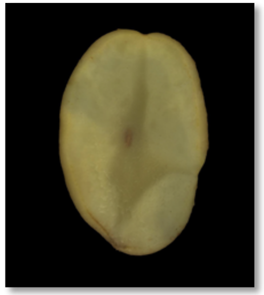 | Hollow hearts | Others (89.18%) |
| 192_1_10_masked | 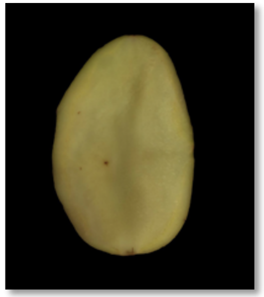 | Others | Hollow hearts (93.06%) |
| 23_1_9_masked | 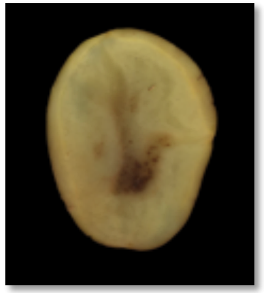 | Others | Hollow hearts (98.70%) |
| 160_2_2_masked | 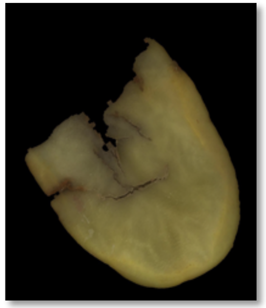 | Others | Hollow hearts (99.20%) |
